# CompScore: boosting structure-based virtual screening performance by incorporating docking scoring functions components into consensus scoring

**DOI:** 10.1101/550590

**Authors:** Yunierkis Perez-Castillo, Stellamaris Sotomayor-Burneo, Karina Jimenes-Vargas, Mario Gonzalez-Rodriguez, Maykel Cruz-Monteagudo, Vinicio Armijos-Jaramillo, M. Natália D. S. Cordeiro, Fernanda Borges, Aminael Sánchez-Rodríguez, Eduardo Tejera

## Abstract

Consensus scoring has become a commonly used strategy within structure-based virtual screening (VS) workflows with improved performance compared to those based in a single scoring function. However, no research has been devoted to analyze the worth of docking scoring functions components in consensus scoring. We implemented and tested a method that incorporates docking scoring functions components into the setting of high performance VS workflows. This method uses genetic algorithms for finding the combination of scoring components that maximizes the VS enrichment for any target. Our methodology was validated using a dataset that contains ligands and decoys for 102 targets that has been widely used in VS validation studies. Results show that our approach outperforms other methods for all targets. It also boosts the initial enrichment performance of the traditional use of whole scoring functions in consensus scoring by an average of 45%. Finally, our methodology was shown to be outstandingly predictive when challenged to rescore external (previously unseen) data. CompScore is freely available at: http://bioquimio.udla.edu.ec/compscore/.

## 1. Introduction

Structure-Based Drug Discovery (SBDD) uses the 3D-structure of proteins (targets) and compounds (ligands) in order to predict their potential interaction. Among SBDD applications, virtual screening (VS) aims at, given a large database of chemical compounds, rank them from highest to lowest probabilities of binding to a target of interest (Romano, 2007). One of the most widely used tool in SBDD is molecular docking which has been particularly useful in VS (Lionta *et al.*, 2014).

Any docking method has two main components: the conformational exploration of the receptor-ligand complex and the evaluation of the predicted interactions in the complex. Among them, scoring functions remain the weakest component. Deficiencies in the scoring functions can be explained by the complexity in estimating the binding energy between the protein and the ligand (Drwal and Griffith, 2013). Each scoring function uses different physicochemical descriptions and parameters to estimate the binding affinity of the complex. However, no individual scoring function accounts for all the physicochemical events that are involved in the protein-ligand interactions and the computational estimation of the binding energy is just an approximation to the reality (Kitchen *et al.*, 2004; Yunta, 2016). For example, one scoring function might be very good at treating solvation effects but not at taking into account shape complementarity. To overcome these deficiencies, consensus scoring (CS) has emerged as a strategy that has shown to outperform single scoring functions since it combines information from a variety of them and compensates their individual weaknesses (Wang and Wang, 2001; Houston and Walkinshaw, 2013; Klingler *et al.*, 2015; Ballester *et al.*, 2012).

CS studies differ in the target, the combination of scoring functions they used and also in the method used to aggregate them. There are many reports of approaches of variable complexity to fuse scoring functions. Some of them employ classical aggregation methods such as majority voting, rank, maximum, intersection and minimum (Yang *et al.*, 2005; Plewczynski *et al.*, 2011). Some other methods include multilinear regression, non-linear regression and multivariate analysis (Feher, 2006).

In this sense, the use of machine learning (ML) approaches for the design of CS strategies has shown promising results and proved to be an innovative and effective way to overcome difficulties in structure-based CS (James *et al.*, 2009). Consequently, it leads to more accurate and general outcomes (Ballester and Mitchell, 2010). In a recent study, an unsupervised machine learning algorithm for structure-based VS (Gradient Boosting) was proposed (Ericksen *et al.*, 2017). This method was tested on 21 targets from DUD-E database (Mysinger *et al.*, 2012) and that the ML-CS strategies obtained better results in comparison to the traditional CS and the individual scoring methods. A supervised CS strategy using Random Forest to successfully predict protein-ligand binding poses has also been proposed (Teramoto and Fukunishi, 2007).

So far, the existing CS strategies consist in combining the scoring functions from one or more software employing different methods. To the best of our knowledge, none of them has focus on the potential of scoring functions components for CS. Furthermore, many of these studies develop general models for any problem and have been validated on a limited number of cases study. Within this panorama we aim at addressing one question: Can individual scoring functions components be more effective than whole scoring functions for CS?

Here, we propose the CompScore algorithm as a universal tool for CS that exploits the information provided by the components of scoring functions. In CompScore the scoring functions are decomposed into their components and a genetic algorithm is used to find the combination of them that maximizes the VS enrichment in each case study. This approach leads to tailored CS schemes for every target. Our methodology is extensively validated and found to be superior to any other tested scoring approach. CompScore is freely available through a Web Service at: http://bioquimio.udla.edu.ec/compscore/

## 2. Methods

### 2.1 Datasets and preparation

Validation datasets were downloaded from the Directory of Useful Decoys, Enhanced (DUD-E) (Mysinger *et al.*, 2012). All the 102 targets from the DUD-E database were selected for the validation of our methodology. We employed the same receptors structures and docking boxes as in the DUD-E. Receptors preparation included the addition of hydrogen atoms and partial atomic charges and their conversion to MOL2 format. Receptors were prepared with UCSF Chimera (Pettersen *et al.*, 2004).

In the DUD-E validation, docking calculations were performed with DOCK 3.6 (Mysinger and Shoichet, 2010). The DUD-E database contains more than one scored conformation per compound. Given that in a regular large scale VS campaign only one conformation per compound is analyzed, we filtered the provided conformers to keep the top-scored one of each compound. Ligands were converted to MOL2 format using UCSF Chimera. Atomic partial charges for compounds provided in the DUD-E were preserved. A summary of the composition of the dataset for each target is provided as Supplementary Data in Table TS1.

### 2.2 Rescoring

Conformers were rescored with DOCK 6.8 (Allen *et al.*, 2015), OEDocking (Kelley *et al.*, 2015) and Gold (Jones *et al.*, 1997) using each scoring function default parameters. Python scripts were created to rescore molecules using OEDocking Toolkits and to produce scoring tables summaries without structures output. A total of 15 scoring functions were computed with these software and they are listed in Table 1. The Dock 3.6 scoring values provided in the DUD-E were included in our calculations.

**Table 1.**
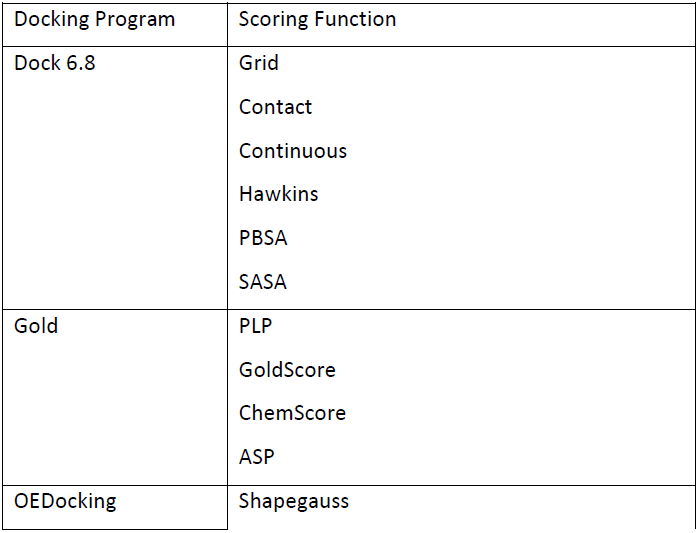
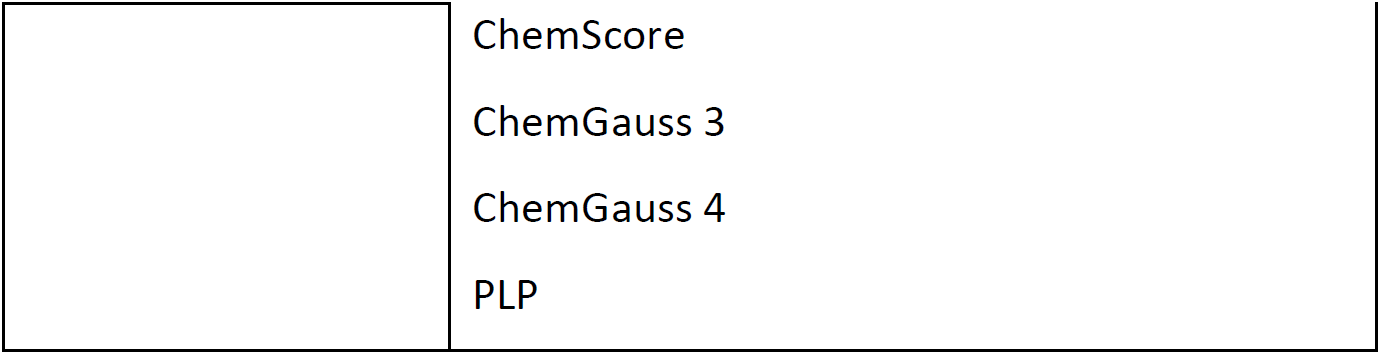
Scoring functions computed per docking program

### 2.3 The CompScore algorithm

The CompScore algorithm searches for the combination of scoring functions components that maximizes a pre-selected VS enrichment metric. In this study, 15 scoring functions obtained by three docking programs were computed for each compound in the DUD-E. With this amount of scoring functions, the exhaustive search of all possible combinations of scoring functions (of size 1 to 16, 65535 combinations) can be completed in a short time. However, these 15 scoring functions account for 87 scoring components in total. This leads to a combinatorial explosion making impossible to evaluate the VS quality of all of their possible combinations. For this reason, the CompScore algorithm implements a genetic algorithm (GA) to search for the combination of scoring components that maximizes the desired VS enrichment metric.

#### 2.3.1 Input and output data

A helper python script was developed to summarize the rescoring results in a data table. This script extracts the molecules’ scoring information, including the value of scoring function components, from DOCK 6.8 scoring files, OEDocking scoring tables and Gold log files. It currently supports the scoring functions listed in Table 1 and is freely available for download from the CompScore server. In addition to scoring information, the scoring data table contains an ID, the number of heavy atoms and a classification as either ligand (1) or decoy (0) for each molecule. Compounds for which any scoring function failed were assigned score values extreme enough to be ranked at the end of the function’s ranked list.

The output of the algorithm is a log file containing information relative to which scoring functions must be combined to maximize the desired VS enrichment metric and enrichment values. A second output file with a ranked list of the input compounds with aggregated scores is provided as output. For more information on data input and output for the CompScore algorithm, see the help pages available at http://bioquimio.udla.edu.ec/compscore-help/.

#### 2.3.2 Scoring components pre-processing

The first step of the CompScore algorithm is to convert the scores to relative rankings. For this, samples are ranked according to each scoring function and the relative ranks are computed as the ranks divided by the number of compounds in the dataset. For ranking molecules we considered, from the definition of the scoring functions within the docking programs, whether scores should be ranked in descending or in ascending order.

To avoid redundancy in the input features, the correlation between the rankings produced by the scoring functions components are analyzed and only one ranking among those having a correlation above certain threshold are kept on the dataset. From a set of correlated rankings, the one appearing first in the dataset is removed. In our validations, the allowed correlation between relative rankings was set to 0.95. In addition, it is possible to establish a minimum number of scores levels that a scoring function component must provide. For example, a scoring component with the same value for all samples will produce a meaningless (constant) ranking of the compounds. The same applies to scoring functions components providing very few different values (levels). The CompScore algorithm provides the possibility of excluding scoring function components having a few levels. For our calculations, the scoring components providing less than four different score levels were excluded.

#### 2.3.3 Virtual Screening enrichment metrics

VS protocols validation aims at obtaining the highest enrichment of ligand molecules at the beginning of the ranked list. This is achieved through the maximization of an enrichment metric, which is selected depending on the objective of the VS campaign. In the CompScore algorithm, either the Enrichment Factor (EF) or BEDROC metrics can maximized. These metrics have been extensively used for the estimation of the enrichment capacity of VS workflows, see (Truchon and Bayly, 2007; Huang *et al.*, 2006; Perez-Castillo *et al.*, 2018; Helguera *et al.*, 2016) for definitions and applications. The main difference between them is that EF measures the enrichment of ligands at a specific fraction of the ranked list while BEDROC accounts for early enrichment. For the maximization of EF, the fraction of screened data at which this metric is expected to be maximum must be provided. On the other hand, if BEDROC is to be maximized, the value of the α parameter for BEDROC calculation should be provided as input.

#### 2.3.4 GA-guided consensus scoring

At this point, scoring components have been converted to relative rankings and filtered as described in section 2.3.2. Also, an enrichment metric has been selected for maximization (2.3.3). Now, the CompScore algorithm searches for the combination of scoring functions components providing the highest enrichment. The GA-guided search takes place in the context of a feature selection problem that maximizes an objective function and was implemented using the DEAP framework in Python (Fortin *et al.*, 2012).

Individuals in the population for GA evolution were coded as binary vectors of length equals to the number of input scoring functions components. Bits set to 1 indicate that the corresponding scoring component is considered for CS, while those corresponding to 0-coded bits are excluded. In our validations we used 100 individuals in the population and they evolved for 1000 generations. The initial population was randomly created. The selection operator was set to a tournament of size 2. The two points crossover operator was used for crossover and mutation proceeded according to the bit flipping operator.

The objective function of the GA was set to either the EF or BEDROC metrics. To obtain the ranked list of compounds for enrichment computation, a subset of relative rankings was aggregated using the arithmetic mean and sorted in ascending order. Then, the chosen enrichment metric was computed for the aggregated ranking of compounds.

After finishing the GA evolution, the best solution was chosen as the individual in the population with the highest fitting (VS enrichment metric). A bootstrap cross-validation was performed to the best solution to evaluate its robustness. For this, 1000 bootstrap samples containing the same number of ligands and decoys as the whole target dataset were generated. All calculations were performed using a single core in a computer equipped with two Intel Xeon CPU E5-2690 and with 128 GB of RAM. The program was developed with Python 3.6 installed within the Anaconda Distribution and package management system.

### 2.4 External validation

For external validation, targets’ datasets were randomly split into training and external validation sets containing 80% and 20% of data, respectively. To maintain the same proportion of ligands and decoys in these sets as in the whole datasets, splitting was performed for ligands and decoys separately. That is, the training sets contained 80% of ligands and 80% of decoys and the external validation sets contained the remaining compounds. Then, we used the training data to obtain the set of scoring functions components maximizing the initial enrichment of actives by employing BEDROC with α=160.9 as fitness function. Afterward, the resulting CompScore model was used to rescore the compounds on the external dataset and initial enrichment was measured over this rescored data. This external validation procedure was repeated 100 times for each target.

## 3. Results and discussion

We performed calculations with the aim of establishing the VS performance of the CompScore algorithm and testing how it compares to that of other scoring schemes. To this end, we explored all possible combinations of the 16 scoring functions of size 1 to 16, totaling 65535 CS solutions. This experiment provides the maximum possible enrichment value that could be achieved by combining any of the 16 scoring functions and is referred to as Exhaustive Search (ES). The best performing scoring function component (including whole scoring functions) was also analyzed and is referred to as Best Individual Scoring Component (BISC). The fourth CS approach consisted in the aggregation of all non-correlated and non-close to constant scoring components. This last approach is referred to as All Scoring Components (ASC) from here on. Finally, we explored the worth of incorporating the values of the scoring components weighted by the number of heavy atoms into the CS procedure.

As previously mentioned, the CompScore algorithm can be used to find the combination of scoring functions components that maximizes either EF or BEDROC employing any metric specific parameter. Although both metrics evaluate the enrichment ability of VS methodologies, unlike EF, BEDROC is able to account for the early enrichment factor. Thus, we made experiments with both metrics. For the sake of simplicity, from here on all analyses will focus on BEDROC computed for the α parameter equals to 160.9 and on EF computed for the first 1% of screened data. In contrast to BEDROC, that is bounded between 0 and 1, EF is an unbounded metric which makes difficult its use in comparisons between the VS performance scoring method across different datasets. For this reason, the maximum possible EF (for a perfect ranking) was computed for each dataset and comparative analyses were carried out with the fraction of this optimal EF achieved by the VS scoring methods.

The results obtained for all the DUD-E targets employing the four CS strategies previously described are provided as Supplementary Data. Table TS2 contains the enrichment metrics obtained with the different scoring approaches when BEDROC with α=160.9 is used as the model selection criterion. On the other hand, table TS3 includes the equivalent information when EF at the first 1% of scoring data is employed as criterion for the selection of the best VS strategy. The results obtained for the early enrichment performance of the four CS strategies explored are summarized in Fig 1. In addition to the best solution per scoring strategy, we also summarize the best performance obtained per target with any of the ASC, BISC and ES strategies (cyan).

**Fig 1.**
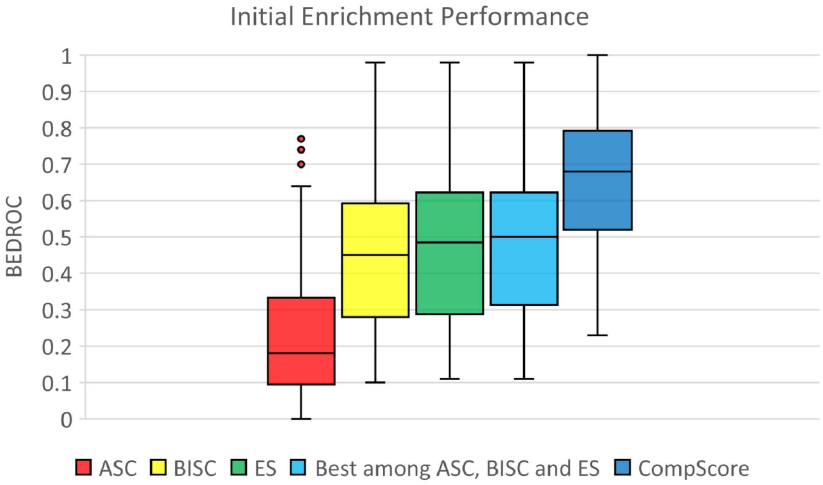
Box plot of the Initial enrichment performance of the evaluated scoring strategies. Initial enrichment is measured as BEDROC for α=160.9. Black lines within the boxes indicate the value of the median. Plot is built with the values of BEDROC for all DUD-E targets as provided in Supplementary Data table TS2.

Fig 1 shows that the CompScore methodology (blue) outperforms the ASC, BISC and ES strategies and that it is the approach that provides the highest value of minimum early enrichment. The improvement resulted statistically significant (p=8·10^-8^) according to the two-tail unpaired t-Test. While any of the methodologies used for comparison (cyan) yield BEDROC values around 0.1, the minimum value obtained with the CompScore methodology is 0.23. Furthermore, more than 50% of the top BEDROC values obtained with CompScore are higher than the top 25% values provided by any of the other methodologies. More important, when a target by target analysis is performed, it is observed that the proposed methodology outperforms any of the other methods for all of them (see Supplementary Data table TS2).

The worst initial enrichment is obtained when all scoring components are aggregated (ASC). This result is expected since no scoring components selection is performed before aggregation. It is well established that any consensus decision making system must contain a diverse subset of decision makers which are meaningful for the problem under investigation (Polikar, 2006). The latter is addressed by the CompScore algorithm through the GA search.

Another interesting result is that the performances of the BISC and ES approaches are highly similar. Thus, in general it can be concluded that a single scoring component can achieve a similar VS performance as the best combination of whole scoring functions. A deeper analysis of the data presented in Supplementary Data table TS2 shows that for 12 targets a single scoring function component (BISC approach) produces BEDROC values more than 5% higher than the best combination of whole scoring functions derived from the ES approach. This improvement can reach up to 73.5% for the mmp13 target. This finding supports our hypothesis that docking scoring functions components provide meaningful information for implementing VS workflows and that in some cases they can be more relevant than whole scoring functions.

The execution time analysis shows that, in average, the CompScore algorithm requires 356 seconds while 150 seconds are required for completing the exhaustive exploration of the combinations of the 16 scoring functions. It must be considered that, according to the run time for the ES approach with 16 scoring functions, for completing the exhaustive exploration of 18 scoring functions the estimated required time for the exhaustive exploration of all their possible combinations will almost double that needed by CompScore. Moreover, the bootstrap cross-validation of our methodology shows that it is stable relative to changes in the dataset composition (see Supplementary Data table TS2).

We also analyzed the BEDROC increase achieved by our methodology across all targets relative to the other tested scoring methodologies. This analysis reveals that CompScore improves, in average, the performance of any other tested method in 45%. However, there are large variations in this improvement among all DUD-E targets as shown in Fig 2. According to Fig 2, for more than 50% of the targets BEDROC improves in more than 33.87% and in more than 64.29% for 25% of the targets when the CompScore methodology is employed.

**Fig 2.**
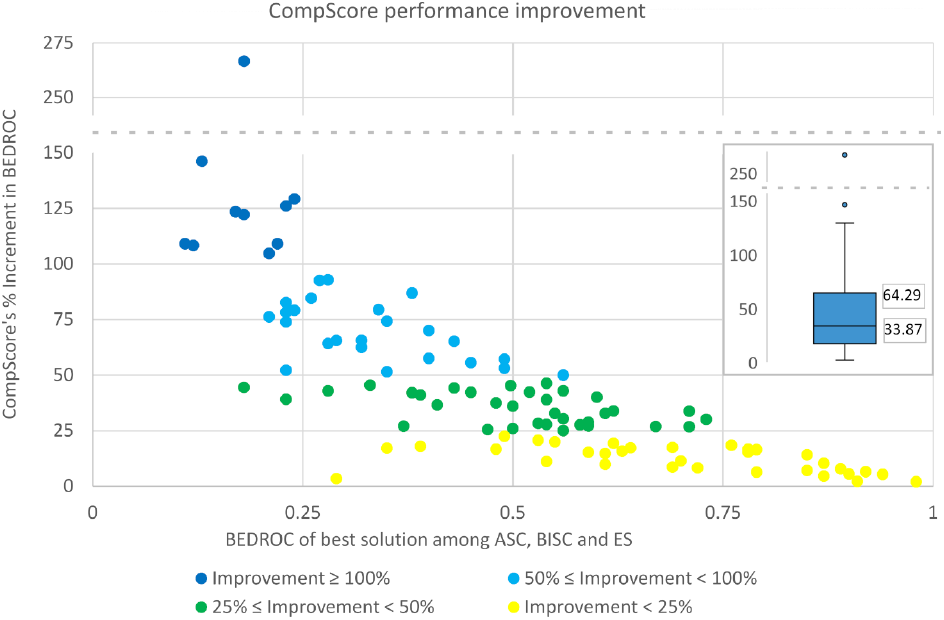
Percent of improvement in BEDROC of the CompScore method relative to the best performing VS models obtained with either the ASC, BISC and ES approaches.

For the 18 targets with BEDROC lower than 0.25 according to all of the ASC, BISC and SE approaches, only two of them increase their BEDROC in less than 50% when CompScore is employed for scoring (see Fig 2). For 10 of them BEDROC increases in more than 100%, that is, BEDROC according to our methodology is more than twice the maximum value obtained with any of the other tested approaches. On the other hand, targets for which BEDROC higher than 0.75 could be obtained with other methods presented an improvement of this metric lower than 25%. The largest improvement in BEDROC is observed for the ampc target with 266.67% (over 3.5 times). For this target the BISC approach yields the Dock 3.6 scoring function as the best performing one with BEDROC=0.16. The next best performing approach is the ES with BEDROC=0.18 through the combination of the scoring functions: Dock 3.6, Dock 6.8 Pbsa, Gold Goldscore and OEDocking ChemGauss 4. On the other side, the CompScore algorithm achieves a value of BEDROC=0.66 by aggregating two scoring functions (Dock 3.6 and OEDocking ShapeGauss) and 12 scoring functions components. These components include four from Dock 6.8, two from Gold and six from OEDocking.

All the solutions found by CompScore include scoring components of at least two different docking software. Specifically, 60 solutions contain scoring functions components from the four docking software, 37 from 3 software and 5 of only two programs. Also, for nine targets: cp3a4, def, fak1, hdac8, kit, kpcb, pgh1, reni and rxra the best solution found with our algorithm contains no whole scoring functions but only their components. Regarding the representativeness of each docking software in the solutions found by CompScore, Dock 3.6 scoring function appears in the best combination for 65 targets, Dock 6.8 scoring components in 101, OEDocking scoring components in 96 and Gold ones in 99. In addition, 17 scoring functions components out of 87 were not part of any CompScore solution because of being either constant or correlated in more than 90% of the targets under investigation. The log files containing the list of the scoring functions included in each target’s best solution are provided at the CompScore’s web site.

Hitherto, all the evidence indicates that decomposing the docking scoring functions into their components and the aggregation of a subset of them ensures high performance VS workflows in terms of initial enrichment. We also evaluated the performance of CompScore when VS campaigns focus on maximizing EF. When the maximum value of EF achieved by each method is analyzed, the results are similar to those obtained with BEDROC as shown in Fig 3. For EF, the proposed algorithm also outperforms the rest of the test scoring schemes. The performance improvement for EF is also statistically significant (p=1.5·10^-7^). As for BEDROC, more than the top 50% performing solutions found by the CompScore methodology are higher than the top 25% solutions found by any of the other methods. The target by target analysis of the achieved fraction of the maximum possible EF yields that for five targets our methodology is unable to improve this metric relative to the other approaches. Also, there is a 3.94% decrease in performance for the mmp13 target relative to the BISC method. Nevertheless, for 50% of the targets the EF improvement is higher than 39.84% and for 25% of them it is of more than 70% relative to any of the other methodologies.

**Fig 3.**
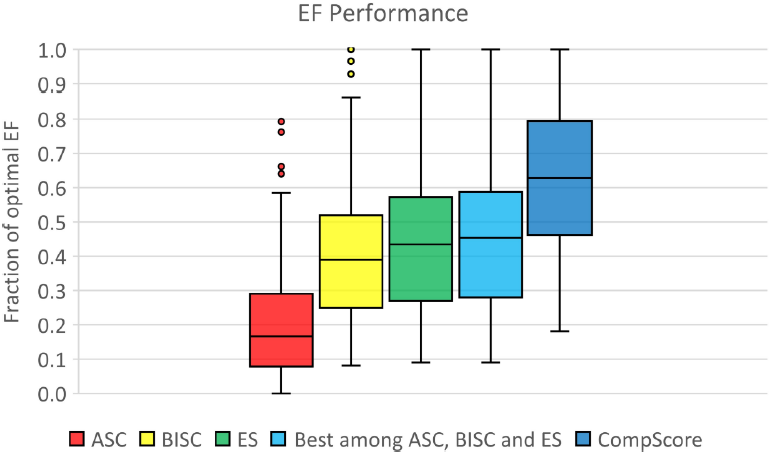
Box plot of the EF performance of the evaluated scoring strategies. EF is measured for a fraction of screened data equals to 0.01. Black lines within the boxes indicate the value of the median. Plot is built with the values of EF for all DUD-E targets as provided in Supplementary Data table TS3.

We explored whether the addition of the docking scoring components weighted by the number of heavy atoms to the GA search could improve the previously presented results. Given that using the weighted scores in the ES approach for the 16 computed scoring functions will take an estimate of more than 100 days per target, this approach was excluded from these analyses. The results of the inclusion of the weighted scores into CompScore are presented as Supplementary Data in tables TS4 and TS5 for BEDROC and EF, respectively, and summarized in Fig 4.

**Fig 4.**
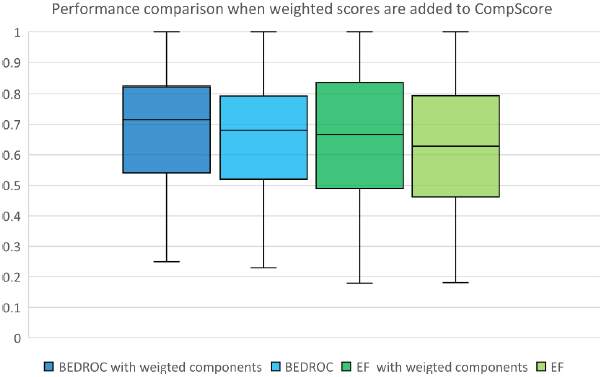
Performance comparison when the weighted scores are added to the CompScore algorithm. EF is represented as the fraction of its maximum possible value for each target.

From Fig 4 it can be observed that the inclusion of the weighted scores dos not increase, in average, the performance of the CompScore algorithm when either BEDROC or EF are used as VS selection metrics. However, a closer look at the results of this last experiment shows that the inclusion of the weighted scoring components can increase in 20% or more the initial enrichment of the VS models of the targets dhi1 (20%), hivint (25%), mcr (25%) and hivpr (50%). Likewise, the EF for targets aldr, mmp13, hivint and hivpr increase by 22%, 22%, 24% and 27%, respectively, when weighted scoring components are considered in the algorithm. In DUD-E, ligands and decoys have similar molecular weight. Thus, we speculate that in databases with uneven distribution of molecules’ sizes the inclusion of the weighted scoring components can translate into improved VS workflows.

The only criterion that can be used to evaluate the generalization capability of CompScore to new data is its performance on a data subset not used for model training. This external validation experiments were performed following the procedure described in the Methods section. External validation results are provided as Supplementary Data in table TS6 and summarized in Fig 5. Results are presented for the average BEDROC over the 100 training/external validation splits performed for each target. First, it can be seen that the initial enrichment obtained for the training data does not differ from that of the previous experiments when all data was used to train CompScore models (blue and cyan boxes, respectively). This result is in agreement with the robustness shown by CompScore in the bootstrap cross-validation experiments.

**Fig 5.**
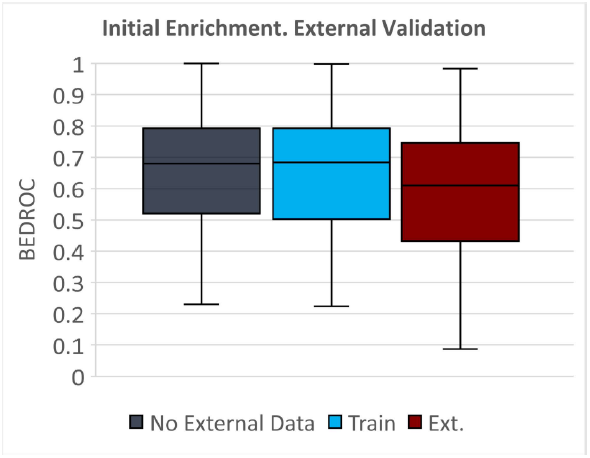
External validation results. CompScore enrichment when all data is used for training is presented as a dark blue box. Cyan and red boxes correspond to the training and external data predictions, respectively

Likewise, the enrichment values observed for the external validation sets (red box in Fig 5) are close to those observed for the training set. The closeness of the enrichment on the external validation set to that of the training set shows that CompScore provides predictive models. The data presented in table TS6 shows that the difference in average BEDROC between the training and external validation sets is lower than 0.1 for 85 out of 102 targets. That is, BEDROC difference between training and prediction is lower than 10% of its theoretical maximum for more that 83% of the DUD-E targets. In addition, BEDROC differences higher than 0.2 can only be observed for the targets mk01, cxcr4 and inha. These targets are among the lowest represented ones in the DUD-E database with 78, 40 and 44 ligands, respectively. The larger differences in BEDROC between the training and external validation sets in these targets could be a consequence of the low amount of data available for training when 20% of it is removed. We consider that these reductions in predictability for these few targets do not represent a loos of generalization in CompScore. In summary, the results of the external validation show that the proposed methodology is not only able to provide high initial enrichment for the training data, but also to extend this enrichment to previously unseen data.

## 4. Conclusions

Here, we introduce CompScore, a simple, fast, interpretable and universal algorithm that incorporates for the first time the idea of decomposing docking scoring functions into their components for consensus scoring in VS. The problem of the combinatorial explosion due the large pool of features that can be extracted from docking scoring functions, is addressed by using a GA as feature selection tool that searches for a subset of them maximizing either the BEDROC or EF metrics. We evaluated our method using the whole DUD-E database and compared its performance with that obtained with all the 65535 combinations of 16 diverse scoring functions from 3 docking software. In addition, we accomplished comparisons with the best performing scoring component and with the aggregation of all of them.

In terms of initial enrichment, for all DUD-E targets CompScore outperformed the rest of that we tested, including the exhaustive exploration of the 16 scoring functions. Our method was also found to highly improve, in more than 100%, the performance for targets for which the rest of the tested methods provided very low initial enrichment. The obtained results also highlight the importance of using a diverse set of scoring functions for consensus scoring since all found solutions included scoring components from at least two docking software. Similar conclusions could be extracted for the VS experiments performed using EF as objective function. It must be highlighted that this exceptional performance could be obtained in less than 8 minutes for more than 75% of the targets. We also showed that the inclusion of the weighted scores can, in some cases, improve the VS performance of consensus scoring strategies. Finally, CompScore was shown to achieve a VS performance on unseen data similar to that observed for the training data. The latter demonstrated that the models proposed by CompScore are able to provide rankings highly enriched with ligands when new collections of compounds are predicted.

Altogether, our results show that scoring functions components are more effective than whole functions for setting high performance VS protocols. In terms of docking calculations time, no extra effort is necessary to apply the CompScore algorithm since all the information that it requires is included in the docking output files provided by the software. Finally, we propose that docking scoring functions breakdown into their components should become a routinary task for the development of CS workflows.

CompScore can be seen as a take the best of each world (docking software and scoring functions) approach. We are aware that the affirmation that the use of scoring components for CS outperforms current CS methods based on whole scoring functions, can be polemic. However, we expect that the availability of CompScore along with the good performance that it achieved will attract the attention of researchers to further corroborate or reject our hypothesis.

## References

Allen, W.J. et al. (2015) DOCK 6: Impact of new features and current docking performance. J. Comput. Chem., 36, 1132–1156.

Ballester, P.J. et al. (2012) Hierarchical virtual screening for the discovery of new molecular scaffolds in antibacterial hit identification. J. R. Soc. Interface, 9, 3196–3207.

Ballester, P.J. and Mitchell, J.B.O. (2010) A machine learning approach to predicting protein–ligand binding affinity with applications to molecular docking. Bioinformatics, 26, 1169–1175.

Drwal, M.N. and Griffith, R. (2013) Combination of ligand- and structure-based methods in virtual screening. Drug Discov. Today Technol., 10, e395–e401.

Ericksen, S.S. et al. (2017) Machine Learning Consensus Scoring Improves Performance Across Targets in Structure-Based Virtual Screening. J. Chem. Inf. Model., 57, 1579–1590.

Feher, M. (2006) Consensus scoring for protein–ligand interactions. Drug Discov. Today, 11, 421–428.

Fortin, F.-A. et al. (2012) DEAP: Evolutionary Algorithms Made Easy. J. Mach. Learn. Res., 13, 2171–2175.

Helguera, A.M. et al. (2016) Ligand-Based Virtual Screening Using Tailored Ensembles: A Prioritization Tool for Dual A2AAdenosine Receptor Antagonists / Monoamine Oxidase B Inhibitors. Curr. Pharm. Des., 22, 3082–3096.

Houston, D.R. and Walkinshaw, M.D. (2013) Consensus Docking: Improving the Reliability of Docking in a Virtual Screening Context. J. Chem. Inf. Model., 53, 384–390.

Huang, N. et al. (2006) Benchmarking Sets for Molecular Docking. J. Med. Chem., 49, 6789–6801.

James, L.M. et al. (2009) Machine Learning in Virtual Screening. Comb. Chem. High Throughput Screen., 12, 332–343.

Jones, G. et al. (1997) Development and validation of a genetic algorithm for flexible docking11Edited by F. E. Cohen. J. Mol. Biol., 267, 727–748.

Kelley, B.P. et al. (2015) POSIT: Flexible Shape-Guided Docking For Pose Prediction. J. Chem. Inf. Model., 55, 1771–1780.

Kitchen, D.B. et al. (2004) Docking and scoring in virtual screening for drug discovery: methods and applications. Nat Rev Drug Discov, 3, 935–949.

Klingler, F.-M. et al. (2015) Probing metallo-β-lactamases with molecular fragments identified by consensus docking. Bioorg. Med. Chem. Lett., 25, 5243–5246.

Lionta, E. et al. (2014) Structure-Based Virtual Screening for Drug Discovery: Principles, Applications and Recent Advances. Curr. Top. Med. Chem., 14, 1923–1938.

Mysinger, M.M. et al. (2012) Directory of Useful Decoys, Enhanced (DUD-E): Better Ligands and Decoys for Better Benchmarking. J. Med. Chem., 55, 6582–6594.

Mysinger, M.M. and Shoichet, B.K. (2010) Rapid Context-Dependent Ligand Desolvation in Molecular Docking. J. Chem. Inf. Model., 50, 1561–1573.

Perez-Castillo, Y. et al. (2018) A desirability-based multi objective approach for the virtual screening discovery of broad-spectrum anti-gastric cancer agents. PloS One, 13, e0192176.

Pettersen, E.F. et al. (2004) UCSF Chimera-a visualization system for exploratory research and analysis. J. Comput. Chem., 25, 1605–1612.

Plewczynski, D. et al. (2011) VoteDock: Consensus docking method for prediction of protein–ligand interactions. J. Comput. Chem., 32, 568–581.

Polikar, R. (2006) Ensemble based systems in decision making. IEEE Circuits Syst. Mag., 6, 21–45.

Romano, T.K. (2007) Structure-Based Drug Design: Docking and Scoring. Curr. Protein Pept. Sci., 8, 312–328.

Teramoto, R. and Fukunishi, H. (2007) Supervised Consensus Scoring for Docking and Virtual Screening. J. Chem. Inf. Model., 47, 526–534.

Truchon, J.-F. and Bayly, C.I. (2007) Evaluating virtual screening methods: good and bad metrics for the ‘early recognition’ problem. J. Chem. Inf. Model., 47, 488–508.

Wang, R. and Wang, S. (2001) How Does Consensus Scoring Work for Virtual Library Screening? An Idealized Computer Experiment. J. Chem. Inf. Comput. Sci., 41, 1422–1426.

Yang, J.-M. et al. (2005) Consensus Scoring Criteria for Improving Enrichment in Virtual Screening. J. Chem. Inf. Model., 45, 1134–1146.

Yunta, M.J.R. (2016) Docking and Ligand Binding Affinity: Uses and Pitfalls. Am. J. Model. Optim., 4, 74–114.

