## Supplementary Data for "CompScore: boosting structure-based virtual screening performance by incorporating docking scoring functions components into consensus scoring"

[8] CIQUP/Departamento de Química e Bioquímica, Facultade de Ciências. Universidade do Porto. Porto 4169-007. Portugal

Table TS4. VS performance of different scoring methods for the DUD-E database when BEDROC with α is set to 160.9 is used as model selection criterion and weighted scoring components are employed 14

Table TS5. VS performance of different scoring methods for the DUD-E database when EF at the first 1% of screened data is used as model selection criterion and weighted scoring components are employed ..... 18

Table TS6. VS performance on external validation experiments. Enrichment is computed as BEDROC with α is set to 160.9. Results are presented as the average BEDROC over the 100 training/external partitions of each target ..... 22

### Table TS1. Composition of the datasets employed in our studies

| Target[a] | Num. Ligands[b] | Num. Decoys[c] | Total molecules[d] |
|---|---|---|---|
| aa2ar | 480 | 31281 | 31761 |
| abl1 | 181 | 10609 | 10790 |
| ace | 281 | 16795 | 17076 |
| aces | 442 | 22384 | 22826 |
| ada | 93 | 5427 | 5520 |
| ada17 | 532 | 35723 | 36255 |
| adrb1 | 247 | 15125 | 15372 |
| adrb2 | 229 | 13230 | 13459 |
| akt1 | 289 | 16048 | 16337 |
| akt2 | 116 | 6770 | 6886 |
| aldr | 159 | 8925 | 9084 |
| ampc | 48 | 2829 | 2877 |
| andr | 222 | 13062 | 13284 |
| aofb | 119 | 6730 | 6849 |
| bace1 | 279 | 18025 | 18304 |
| braf | 152 | 9903 | 10055 |
| cah2 | 491 | 30981 | 31472 |
| casp3 | 199 | 10674 | 10873 |
| cdk2 | 474 | 27757 | 28231 |
| comt | 41 | 3824 | 3865 |
| cp2c9 | 119 | 7246 | 7365 |
| cp3a4 | 170 | 11760 | 11930 |
| csf1r | 166 | 12087 | 12253 |
| cxcr4 | 40 | 3401 | 3441 |
| def | 99 | 5689 | 5788 |
| dhi1 | 330 | 19107 | 19437 |
| dpp4 | 532 | 40776 | 41308 |
| drd3 | 480 | 33359 | 33839 |
| dyr | 229 | 17098 | 17327 |
| egfr | 540 | 34869 | 35409 |
| esr1 | 379 | 20413 | 20792 |
| esr2 | 366 | 20087 | 20453 |
| fa10 | 536 | 28143 | 28679 |
| fa7 | 114 | 6213 | 6327 |
| fabp4 | 47 | 2683 | 2730 |
| fak1 | 99 | 5316 | 5415 |
| fgfr1 | 135 | 7785 | 7920 |
| fkb1a | 111 | 5793 | 5904 |
| fnta | 592 | 51258 | 51850 |

| | | | |
|---|---|---|---|
| fpps | 79 | 8536 | 8615 |
| gcr | 206 | 14146 | 14352 |
| glcm | 54 | 3739 | 3793 |
| gria2 | 157 | 11688 | 11845 |
| grik1 | 100 | 6478 | 6578 |
| hdac2 | 179 | 10155 | 10334 |
| hdac8 | 168 | 10426 | 10594 |
| hivint | 99 | 6631 | 6730 |
| hivpr | 533 | 35564 | 36097 |
| hivrt | 334 | 18293 | 18627 |
| hmdh | 170 | 8726 | 8896 |
| hs90a | 78 | 4822 | 4900 |
| hxk4 | 92 | 4385 | 4477 |
| igf1r | 148 | 9267 | 9415 |
| inha | 44 | 2293 | 2337 |
| ital | 138 | 8459 | 8597 |
| jak2 | 107 | 6450 | 6557 |
| kif11 | 116 | 6832 | 6948 |
| kit | 166 | 10404 | 10570 |
| kith | 57 | 2845 | 2902 |
| kpcb | 133 | 8644 | 8777 |
| lck | 419 | 27279 | 27698 |
| lkha4 | 167 | 7448 | 7615 |
| mapk2 | 100 | 6112 | 6212 |
| mcr | 51 | 4157 | 4208 |
| met | 166 | 11207 | 11373 |
| mk01 | 78 | 4423 | 4501 |
| mk10 | 104 | 6574 | 6678 |
| mk14 | 578 | 35715 | 36293 |
| mmp13 | 572 | 37027 | 37599 |
| mp2k1 | 120 | 8057 | 8177 |
| nos1 | 99 | 8001 | 8100 |
| nram | 98 | 6181 | 6279 |
| pa2ga | 99 | 5139 | 5238 |
| parp1 | 507 | 29928 | 30435 |
| pde5a | 398 | 27437 | 27835 |
| pgh1 | 184 | 10312 | 10496 |
| pgh2 | 415 | 21362 | 21777 |
| plk1 | 106 | 6767 | 6873 |
| pnph | 103 | 6901 | 7004 |
| ppara | 372 | 17807 | 18179 |
| ppard | 240 | 11801 | 12041 |
| pparg | 482 | 25094 | 25576 |

| | | | |
|---|---|---|---|
| prgr | 247 | 14248 | 14495 |
| ptn1 | 130 | 7198 | 7328 |
| pur2 | 50 | 2683 | 2733 |
| pygm | 77 | 3927 | 4004 |
| pyrd | 111 | 6091 | 6202 |
| reni | 104 | 6943 | 7047 |
| rock1 | 100 | 6274 | 6374 |
| rxra | 120 | 6270 | 6390 |
| sahh | 60 | 2821 | 2881 |
| src | 522 | 34184 | 34706 |
| tgfr1 | 133 | 8409 | 8542 |
| thb | 97 | 6375 | 6472 |
| thrb | 456 | 26863 | 27319 |
| try1 | 448 | 25787 | 26235 |
| tryb1 | 148 | 7609 | 7757 |
| tysy | 109 | 6715 | 6824 |
| urok | 162 | 9810 | 9972 |
| vgfr2 | 402 | 22623 | 23025 |
| wee1 | 102 | 6136 | 6238 |
| xiap | 100 | 5130 | 5230 |

[a] Target identification on the DUD-E database (http://dude.docking.org/)
[b] Number of ligands
[c] Number of decoy molecules
[d] Total number of molecules per target

Table TS2. VS performance of different scoring methods for the DUD-E database when BEDROC with α is set to 160.9 is used as model selection criterion

| Target[a] | CompScore | | | | | Exhaustive Search | | | | | Best Individual Scoring Component | | | All Scoring Components |
|---|---|---|---|---|---|---|---|---|---|---|---|---|---|---|
| | Model Size[b] | BEDROC[c] | Boot. BEDROC[d] | Std. Boot. BEDROC[e] | Time (seconds)[f] | Model Size[b] | BEDROC[c] | Boot. BEDROC[d] | Std. Boot. BEDROC[e] | Time (seconds)[f] | BEDROC[c] | Boot. BEDROC[d] | Std. Boot. BEDROC[e] | BEDROC[c] |
| aa2ar | 15 | 0.73 | 0.73 | 0.02 | 1088.92 | 2 | 0.53 | 0.52 | 0.03 | 336.69 | 0.55 | 0.55 | 0.02 | 0.43 |
| abl1 | 10 | 0.6 | 0.6 | 0.04 | 203.64 | 2 | 0.41 | 0.41 | 0.05 | 110.63 | 0.49 | 0.48 | 0.04 | 0.17 |
| ace | 15 | 0.52 | 0.51 | 0.03 | 388.91 | 2 | 0.32 | 0.32 | 0.03 | 172.19 | 0.3 | 0.3 | 0.03 | 0.04 |
| aces | 7 | 0.66 | 0.66 | 0.03 | 481.02 | 3 | 0.55 | 0.55 | 0.03 | 265.62 | 0.46 | 0.46 | 0.03 | 0.13 |
| ada | 14 | 0.75 | 0.74 | 0.04 | 93.63 | 2 | 0.59 | 0.59 | 0.06 | 60.19 | 0.33 | 0.33 | 0.06 | 0.12 |
| ada17 | 5 | 0.78 | 0.78 | 0.02 | 1258.96 | 1 | 0.64 | 0.64 | 0.02 | 430.22 | 0.72 | 0.72 | 0.02 | 0.33 |
| adrb1 | 9 | 0.48 | 0.48 | 0.04 | 349.04 | 5 | 0.26 | 0.26 | 0.03 | 175.22 | 0.16 | 0.16 | 0.03 | 0.16 |
| adrb2 | 12 | 0.42 | 0.42 | 0.04 | 326.43 | 3 | 0.2 | 0.2 | 0.03 | 131.30 | 0.23 | 0.23 | 0.04 | 0.16 |
| akt1 | 17 | 0.68 | 0.68 | 0.03 | 495.25 | 2 | 0.53 | 0.53 | 0.03 | 170.74 | 0.45 | 0.44 | 0.04 | 0.24 |
| akt2 | 12 | 0.64 | 0.63 | 0.05 | 110.44 | 2 | 0.45 | 0.45 | 0.06 | 75.66 | 0.43 | 0.42 | 0.06 | 0.15 |
| aldr | 18 | 0.6 | 0.59 | 0.04 | 148.11 | 1 | 0.54 | 0.53 | 0.04 | 94.55 | 0.54 | 0.53 | 0.05 | 0.07 |
| ampc | 14 | 0.66 | 0.65 | 0.07 | 35.88 | 4 | 0.18 | 0.17 | 0.07 | 35.55 | 0.16 | 0.16 | 0.07 | 0 |
| andr | 13 | 0.54 | 0.54 | 0.04 | 208.22 | 6 | 0.28 | 0.28 | 0.04 | 129.88 | 0.28 | 0.28 | 0.04 | 0.19 |
| aofb | 18 | 0.4 | 0.4 | 0.05 | 107.99 | 8 | 0.28 | 0.28 | 0.05 | 71.74 | 0.21 | 0.21 | 0.05 | 0.06 |
| bace1 | 11 | 0.32 | 0.32 | 0.03 | 523.42 | 4 | 0.23 | 0.23 | 0.03 | 187.28 | 0.23 | 0.23 | 0.03 | 0.16 |
| braf | 8 | 0.61 | 0.61 | 0.04 | 161.23 | 1 | 0.32 | 0.32 | 0.05 | 102.44 | 0.35 | 0.35 | 0.05 | 0.23 |
| cah2 | 22 | 0.52 | 0.52 | 0.03 | 1252.91 | 3 | 0.23 | 0.23 | 0.02 | 374.27 | 0.17 | 0.17 | 0.02 | 0.08 |
| casp3 | 12 | 0.37 | 0.37 | 0.05 | 306.41 | 2 | 0.21 | 0.21 | 0.04 | 107.80 | 0.19 | 0.19 | 0.04 | 0.04 |
| cdk2 | 17 | 0.47 | 0.47 | 0.03 | 760.82 | 6 | 0.37 | 0.37 | 0.03 | 317.24 | 0.34 | 0.34 | 0.03 | 0.23 |
| comt | 21 | 0.99 | 0.99 | 0.01 | 56.58 | 1 | 0.92 | 0.92 | 0.03 | 43.52 | 0.94 | 0.94 | 0.02 | 0.45 |
| cp2c9 | 17 | 0.25 | 0.25 | 0.05 | 121.45 | 6 | 0.12 | 0.12 | 0.04 | 76.81 | 0.11 | 0.11 | 0.04 | 0.06 |
| cp3a4 | 16 | 0.26 | 0.26 | 0.04 | 275.18 | 3 | 0.18 | 0.17 | 0.04 | 115.98 | 0.16 | 0.17 | 0.04 | 0.07 |
| csf1r | 9 | 0.53 | 0.53 | 0.04 | 238.6 | 1 | 0.35 | 0.35 | 0.04 | 121.08 | 0.35 | 0.34 | 0.04 | 0.12 |

| | | | | | | | | | | | | | | |
|---|---|---|---|---|---|---|---|---|---|---|---|---|---|---|
| cxcr4 | 19 | 0.71 | 0.7 | 0.07 | 45.11 | 2 | 0.43 | 0.42 | 0.08 | 40.11 | 0.3 | 0.3 | 0.08 | 0.15 |
| def | 15 | 0.91 | 0.91 | 0.02 | 93.7 | 1 | 0.67 | 0.67 | 0.05 | 60.94 | 0.78 | 0.78 | 0.04 | 0.19 |
| dhi1 | 15 | 0.35 | 0.35 | 0.03 | 359.7 | 3 | 0.23 | 0.23 | 0.03 | 222.89 | 0.19 | 0.19 | 0.03 | 0.05 |
| dpp4 | 10 | 0.64 | 0.64 | 0.02 | 1482.24 | 2 | 0.53 | 0.53 | 0.02 | 437.95 | 0.5 | 0.51 | 0.02 | 0.19 |
| drd3 | 11 | 0.43 | 0.43 | 0.03 | 1093.71 | 5 | 0.24 | 0.24 | 0.02 | 410.68 | 0.17 | 0.17 | 0.02 | 0.04 |
| dyr | 17 | 0.75 | 0.75 | 0.03 | 517.59 | 3 | 0.64 | 0.64 | 0.03 | 190.07 | 0.61 | 0.61 | 0.03 | 0.45 |
| egfr | 10 | 0.75 | 0.75 | 0.02 | 1074.18 | 5 | 0.49 | 0.49 | 0.02 | 392.78 | 0.45 | 0.44 | 0.03 | 0.36 |
| esr1 | 17 | 0.75 | 0.75 | 0.02 | 456.77 | 2 | 0.69 | 0.69 | 0.03 | 219.79 | 0.67 | 0.67 | 0.03 | 0.11 |
| esr2 | 14 | 0.74 | 0.74 | 0.02 | 482.85 | 3 | 0.62 | 0.62 | 0.03 | 229.86 | 0.6 | 0.6 | 0.03 | 0.14 |
| fa10 | 9 | 0.73 | 0.73 | 0.02 | 674.64 | 2 | 0.63 | 0.63 | 0.02 | 310.53 | 0.59 | 0.59 | 0.02 | 0.1 |
| fa7 | 16 | 0.95 | 0.95 | 0.01 | 101.9 | 1 | 0.9 | 0.89 | 0.02 | 69.12 | 0.9 | 0.9 | 0.02 | 0.23 |
| fabp4 | 18 | 0.78 | 0.78 | 0.06 | 35.86 | 1 | 0.7 | 0.69 | 0.07 | 34.33 | 0.7 | 0.69 | 0.07 | 0.1 |
| fak1 | 12 | 0.69 | 0.69 | 0.05 | 100.95 | 4 | 0.54 | 0.54 | 0.06 | 60.17 | 0.5 | 0.49 | 0.07 | 0.27 |
| fgfr1 | 12 | 0.55 | 0.54 | 0.05 | 152.11 | 4 | 0.39 | 0.39 | 0.05 | 83.57 | 0.26 | 0.26 | 0.04 | 0.23 |
| fkb1a | 22 | 0.71 | 0.71 | 0.05 | 106.23 | 7 | 0.38 | 0.38 | 0.06 | 64.28 | 0.34 | 0.34 | 0.06 | 0.14 |
| fnta | 17 | 0.23 | 0.23 | 0.02 | 1631.51 | 2 | 0.11 | 0.11 | 0.01 | 611.25 | 0.1 | 0.1 | 0.01 | 0.05 |
| fpps | 14 | 0.91 | 0.9 | 0.02 | 190.6 | 2 | 0.85 | 0.85 | 0.03 | 85.28 | 0.76 | 0.76 | 0.04 | 0.04 |
| gcr | 15 | 0.48 | 0.47 | 0.04 | 321.36 | 1 | 0.29 | 0.29 | 0.04 | 142.67 | 0.29 | 0.29 | 0.04 | 0.12 |
| glcm | 19 | 0.84 | 0.83 | 0.04 | 59.15 | 2 | 0.6 | 0.61 | 0.07 | 44.55 | 0.49 | 0.48 | 0.07 | 0.24 |
| gria2 | 20 | 0.7 | 0.7 | 0.04 | 318.94 | 2 | 0.61 | 0.61 | 0.04 | 115.11 | 0.59 | 0.59 | 0.04 | 0.48 |
| grik1 | 6 | 0.84 | 0.84 | 0.03 | 107.8 | 2 | 0.56 | 0.55 | 0.06 | 68.70 | 0.55 | 0.55 | 0.05 | 0.32 |
| hdac2 | 14 | 0.74 | 0.74 | 0.03 | 240.08 | 3 | 0.52 | 0.52 | 0.04 | 104.52 | 0.45 | 0.45 | 0.04 | 0.37 |
| hdac8 | 15 | 0.9 | 0.9 | 0.02 | 204.11 | 6 | 0.71 | 0.71 | 0.03 | 105.28 | 0.52 | 0.52 | 0.04 | 0.59 |
| hivint | 13 | 0.32 | 0.31 | 0.06 | 136.17 | 3 | 0.13 | 0.13 | 0.04 | 69.74 | 0.1 | 0.1 | 0.04 | 0.02 |
| hivpr | 12 | 0.3 | 0.3 | 0.02 | 1025.82 | 4 | 0.21 | 0.21 | 0.02 | 378.06 | 0.29 | 0.29 | 0.02 | 0.02 |
| hivrt | 18 | 0.43 | 0.43 | 0.03 | 554.53 | 3 | 0.2 | 0.2 | 0.03 | 207.14 | 0.18 | 0.18 | 0.03 | 0.21 |
| hmdh | 14 | 0.83 | 0.83 | 0.02 | 169.42 | 2 | 0.62 | 0.62 | 0.04 | 89.82 | 0.48 | 0.48 | 0.05 | 0.11 |
| hs90a | 19 | 0.77 | 0.77 | 0.04 | 87.98 | 6 | 0.49 | 0.48 | 0.07 | 53.58 | 0.27 | 0.27 | 0.05 | 0.42 |
| hxk4 | 11 | 0.74 | 0.74 | 0.05 | 75.37 | 2 | 0.58 | 0.57 | 0.06 | 53.26 | 0.34 | 0.35 | 0.07 | 0.51 |
| igf1r | 20 | 0.54 | 0.54 | 0.05 | 209.16 | 3 | 0.38 | 0.38 | 0.05 | 103.68 | 0.34 | 0.34 | 0.05 | 0.23 |
| inha | 18 | 0.7 | 0.69 | 0.07 | 33.37 | 1 | 0.45 | 0.44 | 0.1 | 30.51 | 0.45 | 0.44 | 0.1 | 0.19 |
| ital | 7 | 0.46 | 0.46 | 0.05 | 161.09 | 3 | 0.39 | 0.39 | 0.05 | 90.02 | 0.28 | 0.28 | 0.05 | 0.06 |

| | | | | | | | | | | | | | | |
|---|---|---|---|---|---|---|---|---|---|---|---|---|---|---|
| jak2 | 14 | 0.8 | 0.8 | 0.04 | 117.05 | 3 | 0.56 | 0.56 | 0.05 | 68.63 | 0.51 | 0.51 | 0.06 | 0.37 |
| kif11 | 15 | 0.84 | 0.83 | 0.03 | 130.06 | 4 | 0.79 | 0.79 | 0.03 | 77.14 | 0.77 | 0.77 | 0.04 | 0.05 |
| kit | 9 | 0.38 | 0.38 | 0.04 | 185.31 | 5 | 0.15 | 0.15 | 0.04 | 105.55 | 0.17 | 0.17 | 0.04 | 0.08 |
| kith | 14 | 0.96 | 0.96 | 0.01 | 40.43 | 2 | 0.89 | 0.89 | 0.03 | 35.29 | 0.88 | 0.88 | 0.03 | 0.62 |
| kpcb | 11 | 0.74 | 0.74 | 0.04 | 168.56 | 1 | 0.62 | 0.62 | 0.04 | 92.39 | 0.62 | 0.62 | 0.05 | 0.52 |
| lck | 10 | 0.61 | 0.61 | 0.02 | 993.33 | 2 | 0.34 | 0.34 | 0.03 | 314.28 | 0.31 | 0.31 | 0.03 | 0.31 |
| lkha4 | 17 | 0.59 | 0.59 | 0.05 | 160.92 | 3 | 0.47 | 0.47 | 0.05 | 79.46 | 0.43 | 0.43 | 0.05 | 0.17 |
| mapk2 | 21 | 0.92 | 0.91 | 0.02 | 134.9 | 5 | 0.79 | 0.79 | 0.03 | 65.26 | 0.7 | 0.7 | 0.04 | 0.64 |
| mcr | 18 | 0.52 | 0.52 | 0.08 | 50.35 | 3 | 0.27 | 0.27 | 0.08 | 48.13 | 0.25 | 0.25 | 0.08 | 0.12 |
| met | 10 | 0.67 | 0.67 | 0.04 | 215.84 | 4 | 0.5 | 0.5 | 0.05 | 118.48 | 0.61 | 0.61 | 0.04 | 0.13 |
| mk01 | 19 | 0.63 | 0.62 | 0.06 | 126.09 | 3 | 0.5 | 0.49 | 0.07 | 50.39 | 0.34 | 0.34 | 0.06 | 0.26 |
| mk10 | 19 | 0.56 | 0.55 | 0.05 | 90.35 | 5 | 0.41 | 0.41 | 0.06 | 71.09 | 0.31 | 0.31 | 0.06 | 0.23 |
| mk14 | 14 | 0.41 | 0.41 | 0.02 | 1329.56 | 5 | 0.23 | 0.23 | 0.02 | 442.60 | 0.35 | 0.35 | 0.02 | 0.06 |
| mmp13 | 8 | 0.68 | 0.68 | 0.02 | 1020.82 | 1 | 0.34 | 0.34 | 0.02 | 414.15 | 0.59 | 0.6 | 0.02 | 0.11 |
| mp2k1 | 20 | 0.4 | 0.39 | 0.05 | 158.97 | 3 | 0.18 | 0.18 | 0.04 | 83.43 | 0.12 | 0.12 | 0.04 | 0.16 |
| nos1 | 13 | 0.68 | 0.68 | 0.04 | 142.88 | 6 | 0.43 | 0.43 | 0.05 | 84.22 | 0.5 | 0.5 | 0.05 | 0.39 |
| nram | 16 | 0.95 | 0.95 | 0.01 | 119.16 | 3 | 0.73 | 0.73 | 0.04 | 66.72 | 0.47 | 0.47 | 0.05 | 0.24 |
| pa2ga | 15 | 0.73 | 0.72 | 0.05 | 83.15 | 2 | 0.56 | 0.55 | 0.06 | 57.22 | 0.45 | 0.45 | 0.06 | 0.04 |
| parp1 | 17 | 0.9 | 0.9 | 0.01 | 812.95 | 2 | 0.76 | 0.76 | 0.02 | 356.11 | 0.71 | 0.71 | 0.02 | 0.7 |
| pde5a | 13 | 0.56 | 0.56 | 0.03 | 1007.22 | 2 | 0.48 | 0.48 | 0.03 | 297.26 | 0.47 | 0.47 | 0.03 | 0.29 |
| pgh1 | 12 | 0.4 | 0.4 | 0.05 | 149.71 | 4 | 0.23 | 0.23 | 0.04 | 107.03 | 0.2 | 0.2 | 0.04 | 0.06 |
| pgh2 | 15 | 0.63 | 0.63 | 0.03 | 687.72 | 7 | 0.4 | 0.4 | 0.03 | 233.66 | 0.33 | 0.33 | 0.03 | 0.32 |
| plk1 | 20 | 0.7 | 0.7 | 0.04 | 126.04 | 3 | 0.56 | 0.56 | 0.05 | 70.85 | 0.49 | 0.49 | 0.05 | 0.31 |
| pnph | 23 | 0.9 | 0.9 | 0.02 | 135.09 | 2 | 0.78 | 0.77 | 0.04 | 76.44 | 0.76 | 0.75 | 0.04 | 0.74 |
| ppara | 10 | 0.41 | 0.42 | 0.03 | 369.56 | 4 | 0.23 | 0.23 | 0.03 | 191.24 | 0.14 | 0.14 | 0.02 | 0.07 |
| ppard | 8 | 0.46 | 0.46 | 0.04 | 231.32 | 1 | 0.28 | 0.28 | 0.04 | 117.60 | 0.28 | 0.28 | 0.03 | 0 |
| pparg | 14 | 0.46 | 0.46 | 0.03 | 922.63 | 1 | 0.22 | 0.22 | 0.03 | 274.90 | 0.22 | 0.22 | 0.03 | 0.07 |
| prgr | 21 | 0.75 | 0.75 | 0.03 | 416.19 | 8 | 0.39 | 0.39 | 0.04 | 154.05 | 0.38 | 0.38 | 0.04 | 0.54 |
| ptn1 | 14 | 0.7 | 0.7 | 0.04 | 123.23 | 3 | 0.56 | 0.56 | 0.05 | 82.72 | 0.49 | 0.49 | 0.05 | 0.18 |
| pur2 | 16 | 0.97 | 0.97 | 0.01 | 38.66 | 2 | 0.84 | 0.83 | 0.04 | 33.99 | 0.85 | 0.85 | 0.04 | 0.05 |
| pygm | 9 | 0.55 | 0.55 | 0.07 | 57.72 | 3 | 0.24 | 0.24 | 0.07 | 44.94 | 0.22 | 0.23 | 0.06 | 0.01 |
| pyrd | 20 | 0.81 | 0.81 | 0.03 | 107.26 | 6 | 0.56 | 0.56 | 0.05 | 70.84 | 0.61 | 0.61 | 0.05 | 0.22 |

| | | | | | | | | | | | | | | |
|---|---|---|---|---|---|---|---|---|---|---|---|---|---|---|
| reni | 7 | 0.66 | 0.66 | 0.04 | 143.7 | 3 | 0.48 | 0.48 | 0.06 | 74.09 | 0.38 | 0.37 | 0.06 | 0.23 |
| rock1 | 14 | 0.68 | 0.68 | 0.05 | 100.06 | 3 | 0.4 | 0.39 | 0.06 | 67.75 | 0.29 | 0.29 | 0.05 | 0.34 |
| rxra | 12 | 0.95 | 0.95 | 0.01 | 95.02 | 2 | 0.71 | 0.71 | 0.04 | 67.62 | 0.65 | 0.65 | 0.05 | 0.62 |
| sahh | 18 | 0.98 | 0.98 | 0.01 | 36.9 | 3 | 0.91 | 0.91 | 0.03 | 35.39 | 0.92 | 0.92 | 0.02 | 0.77 |
| src | 10 | 0.53 | 0.53 | 0.02 | 889.05 | 3 | 0.32 | 0.32 | 0.02 | 421.45 | 0.26 | 0.26 | 0.02 | 0.18 |
| tgfr1 | 17 | 0.85 | 0.84 | 0.03 | 182.29 | 5 | 0.67 | 0.67 | 0.04 | 87.56 | 0.58 | 0.57 | 0.05 | 0.45 |
| thb | 15 | 0.81 | 0.81 | 0.03 | 104.52 | 2 | 0.69 | 0.69 | 0.05 | 68.33 | 0.6 | 0.6 | 0.06 | 0.4 |
| thrb | 9 | 0.76 | 0.76 | 0.02 | 946.82 | 3 | 0.59 | 0.59 | 0.02 | 310.40 | 0.49 | 0.49 | 0.03 | 0.12 |
| try1 | 11 | 0.91 | 0.91 | 0.01 | 695 | 2 | 0.87 | 0.87 | 0.01 | 269.52 | 0.83 | 0.83 | 0.02 | 0.12 |
| tryb1 | 13 | 0.79 | 0.79 | 0.03 | 140.13 | 1 | 0.54 | 0.54 | 0.05 | 81.62 | 0.54 | 0.54 | 0.05 | 0.13 |
| tysy | 19 | 0.62 | 0.62 | 0.05 | 116.44 | 2 | 0.43 | 0.42 | 0.06 | 72.73 | 0.38 | 0.38 | 0.06 | 0.28 |
| urok | 13 | 0.93 | 0.93 | 0.01 | 253.93 | 2 | 0.91 | 0.91 | 0.02 | 108.93 | 0.89 | 0.89 | 0.02 | 0.56 |
| vgfr2 | 13 | 0.48 | 0.48 | 0.03 | 708.5 | 1 | 0.32 | 0.32 | 0.03 | 238.85 | 0.33 | 0.33 | 0.03 | 0.13 |
| wee1 | 18 | 1 | 0.99 | 0 | 116.24 | 1 | 0.98 | 0.98 | 0.01 | 64.12 | 0.98 | 0.98 | 0.01 | 0.75 |
| xiap | 14 | 0.96 | 0.96 | 0.01 | 74.34 | 2 | 0.87 | 0.87 | 0.03 | 57.44 | 0.87 | 0.87 | 0.03 | 0.52 |
| Mean | 14.30 | 0.66 | 0.66 | 0.03 | 356.40 | 2.99 | 0.48 | 0.48 | 0.04 | 149.99 | 0.45 | 0.45 | 0.04 | 0.24 |

[(a)] Target identification on the DUD-E database (http://dude.docking.org/)

[(b)] Number of aggregated scoring functions

[(c)] BEDROC of the best performing VS strategy

[(d)] Mean BEDROC on 1000 bootstrap simulations of the best performing VS strategy

[(e)] Standard deviation of the bootstrap cross-validation procedure

[(f)] Run time for each consensus scoring approach

Table TS3. VS performance of different scoring methods for the DUD-E database when EF at the first 1% of screened data is used as model selection criterion

| Target[a] | Opt. EF[b] | CompScore | | | | | | Exhaustive Search | | | | | | Best Individual Scoring Component | | | | All Scoring Components | |
|---|---|---|---|---|---|---|---|---|---|---|---|---|---|---|---|---|---|---|---|
| | | Size[c] | EF[d] | Boot. EF[e] | Std. Boot. EF[f] | Fract. Opt. EF[g] | Time (seconds) [h] | Size[c] | EF[d] | Boot. EF[e] | Std. Boot. EF[f] | Fract. Opt. EF[g] | Time (seconds) [h] | EF[d] | Boot. EF[e] | Std. Boot. EF[f] | Fract. Opt. EF[g] | EF[d] | Fract. Opt. EF[g] |
| aa2ar | 66.17 | 20 | 46.19 | 45.8 | 1.63 | 0.69 | 951.53 | 1 | 33.08 | 33.14 | 1.71 | 0.50 | 354.85 | 33.71 | 33.89 | 1.67 | 0.51 | 24.14 | 0.36 |
| abl1 | 59.61 | 14 | 35.33 | 34.37 | 3.05 | 0.58 | 214.87 | 1 | 22.08 | 21.74 | 2.72 | 0.37 | 110.86 | 25.39 | 24.98 | 2.51 | 0.43 | 10.49 | 0.18 |
| ace | 60.77 | 14 | 29.5 | 28.97 | 2.26 | 0.48 | 418.39 | 2 | 17.06 | 17.19 | 2.02 | 0.28 | 177.24 | 15.99 | 16.17 | 2.05 | 0.26 | 2.13 | 0.04 |
| aces | 51.64 | 13 | 32.25 | 31.96 | 1.83 | 0.62 | 570.06 | 3 | 25.71 | 25.8 | 1.65 | 0.50 | 243.67 | 20.52 | 20.31 | 1.59 | 0.40 | 6.09 | 0.12 |
| ada | 59.35 | 21 | 45.58 | 43.08 | 3.75 | 0.73 | 114.91 | 2 | 32.86 | 32.44 | 4.36 | 0.55 | 60.43 | 19.08 | 18.68 | 3.45 | 0.32 | 5.3 | 0.09 |
| ada17 | 68.15 | 8 | 51.06 | 50.82 | 1.63 | 0.75 | 937.72 | 1 | 37.92 | 37.75 | 1.74 | 0.56 | 415.39 | 46 | 45.67 | 1.5 | 0.67 | 19.15 | 0.28 |
| adrb1 | 62.23 | 14 | 26.27 | 25.91 | 2.33 | 0.42 | 392.3 | 3 | 14.95 | 14.63 | 2.07 | 0.24 | 164.80 | 9.29 | 9.18 | 1.67 | 0.15 | 8.08 | 0.13 |
| adrb2 | 58.77 | 18 | 23.94 | 23.28 | 2.42 | 0.40 | 329.11 | 4 | 10.88 | 10.48 | 1.83 | 0.19 | 141.97 | 11.75 | 11.49 | 1.93 | 0.20 | 7.84 | 0.13 |
| akt1 | 56.53 | 14 | 37.57 | 37.51 | 2.23 | 0.66 | 413.34 | 2 | 26.89 | 26.81 | 2.11 | 0.48 | 167.90 | 23.44 | 23.28 | 2.07 | 0.41 | 12.41 | 0.22 |
| akt2 | 59.36 | 17 | 36.13 | 34.9 | 3.55 | 0.59 | 136.22 | 4 | 24.95 | 24.66 | 3.3 | 0.42 | 71.50 | 23.23 | 23.06 | 3.73 | 0.39 | 10.32 | 0.17 |
| aldr | 57.13 | 18 | 32.02 | 30.75 | 2.97 | 0.54 | 201.36 | 1 | 27 | 27.72 | 2.93 | 0.47 | 90.45 | 27 | 27.56 | 2.92 | 0.47 | 2.51 | 0.04 |
| ampc | 59.94 | 14 | 39.27 | 37.13 | 5.5 | 0.62 | 45.26 | 2 | 10.33 | 10.74 | 4.25 | 0.17 | 35.09 | 8.27 | 7.59 | 3.7 | 0.14 | 0 | 0.00 |
| andr | 59.84 | 12 | 30.14 | 29.57 | 2.52 | 0.49 | 288.34 | 5 | 17.1 | 16.38 | 2.36 | 0.29 | 136.56 | 13.05 | 13.33 | 2.22 | 0.22 | 9.9 | 0.17 |
| aofb | 57.55 | 20 | 21.69 | 20.61 | 3.09 | 0.36 | 147.9 | 6 | 16.68 | 16.11 | 3.05 | 0.29 | 72.20 | 11.68 | 10.85 | 2.73 | 0.20 | 4.17 | 0.07 |
| bace1 | 65.61 | 15 | 17.47 | 16.95 | 2.05 | 0.26 | 508.84 | 1 | 11.41 | 11.5 | 1.93 | 0.17 | 183.25 | 11.77 | 11.7 | 1.88 | 0.18 | 8.56 | 0.13 |
| braf | 66.15 | 11 | 36.68 | 36.27 | 3.16 | 0.55 | 231.19 | 3 | 17.68 | 17.37 | 2.81 | 0.27 | 101.97 | 19.65 | 19.64 | 2.8 | 0.30 | 11.79 | 0.18 |
| cah2 | 64.1 | 23 | 31.74 | 31.51 | 1.71 | 0.49 | 1024.17 | 6 | 13.23 | 12.79 | 1.39 | 0.21 | 362.64 | 12.41 | 12.12 | 1.4 | 0.19 | 4.68 | 0.07 |
| casp3 | 54.64 | 16 | 17.54 | 17.06 | 2.32 | 0.31 | 272.26 | 2 | 12.03 | 11.93 | 1.99 | 0.22 | 116.46 | 10.03 | 9.87 | 1.91 | 0.18 | 1.5 | 0.03 |
| cdk2 | 59.56 | 21 | 27.36 | 26.92 | 1.69 | 0.45 | 807.19 | 5 | 20.2 | 19.86 | 1.55 | 0.34 | 297.64 | 17.05 | 16.87 | 1.43 | 0.29 | 13.47 | 0.23 |
| comt | 94.27 | 21 | 91.85 | 90.34 | 3.13 | 0.96 | 79.93 | 1 | 79.77 | 78.43 | 4.86 | 0.85 | 45.32 | 79.77 | 78.64 | 4.88 | 0.85 | 36.26 | 0.38 |
| cp2c9 | 61.89 | 25 | 15.05 | 14.09 | 2.91 | 0.23 | 165.67 | 4 | 8.36 | 8.07 | 2.35 | 0.14 | 75.81 | 5.85 | 5.3 | 2.01 | 0.09 | 3.35 | 0.05 |
| cp3a4 | 70.18 | 17 | 14.04 | 13.94 | 2.5 | 0.20 | 294.07 | 4 | 11.11 | 10.47 | 2.27 | 0.16 | 118.99 | 10.53 | 10.24 | 2.24 | 0.15 | 4.68 | 0.07 |
| csf1r | 73.81 | 17 | 33.01 | 32.17 | 3.09 | 0.44 | 303.65 | 1 | 24 | 23.63 | 2.93 | 0.33 | 120.77 | 24 | 23.68 | 2.9 | 0.33 | 7.2 | 0.10 |

| | | | | | | | | | | | | | | | | | | |
|---|---|---|---|---|---|---|---|---|---|---|---|---|---|---|---|---|---|---|
| cxcr4 | 86.02 | 19 | 51.61 | 50.37 | 6.83 | 0.59 | 62.36 | 4 | 29.49 | 26.35 | 6.35 | 0.34 | 40.66 | 19.66 | 19.68 | 6.03 | 0.23 | 12.29 | 0.14 |
| def | 58.46 | 17 | 58.46 | 56.89 | 2.27 | 0.97 | 122.1 | 1 | 35.28 | 35.79 | 4 | 0.60 | 62.90 | 42.34 | 42.17 | 3.47 | 0.72 | 8.06 | 0.14 |
| dhi1 | 58.9 | 14 | 19.94 | 19.51 | 1.88 | 0.33 | 452.78 | 3 | 11.78 | 11.7 | 1.74 | 0.20 | 212.26 | 8.76 | 8.84 | 1.5 | 0.15 | 2.42 | 0.04 |
| dpp4 | 77.65 | 13 | 42.95 | 42.56 | 1.76 | 0.55 | 1172.21 | 2 | 34.13 | 34.17 | 1.81 | 0.44 | 478.41 | 33.38 | 33.44 | 1.74 | 0.43 | 13.5 | 0.17 |
| drd3 | 70.5 | 19 | 26.41 | 26.09 | 1.8 | 0.37 | 964.27 | 5 | 14.97 | 14.66 | 1.52 | 0.21 | 358.36 | 9.77 | 10.11 | 1.3 | 0.14 | 3.33 | 0.05 |
| dyr | 75.66 | 20 | 52.18 | 51.67 | 2.86 | 0.68 | 462.34 | 2 | 40.01 | 39.61 | 2.82 | 0.53 | 171.22 | 39.14 | 39.26 | 2.68 | 0.52 | 26.96 | 0.36 |
| egfr | 65.57 | 10 | 49.32 | 48.83 | 1.54 | 0.74 | 971.57 | 5 | 28.45 | 28.59 | 1.61 | 0.43 | 415.57 | 22.72 | 22.82 | 2.3 | 0.35 | 20.32 | 0.31 |
| esr1 | 54.86 | 17 | 43.26 | 42.5 | 1.61 | 0.77 | 502.85 | 1 | 36.4 | 36.4 | 2.11 | 0.66 | 215.12 | 34.29 | 33.97 | 2.14 | 0.63 | 6.07 | 0.11 |
| esr2 | 55.88 | 11 | 41.71 | 41.64 | 1.84 | 0.75 | 472.16 | 3 | 32.17 | 32.27 | 2.12 | 0.58 | 214.72 | 29.99 | 30.2 | 1.98 | 0.54 | 8.45 | 0.15 |
| fa10 | 53.51 | 14 | 39.34 | 38.95 | 1.68 | 0.73 | 786.81 | 3 | 31.32 | 31.3 | 1.53 | 0.59 | 294.69 | 28.71 | 28.61 | 1.46 | 0.54 | 5.03 | 0.09 |
| fa7 | 55.5 | 17 | 55.5 | 55.47 | 0.25 | 1.00 | 130 | 1 | 55.5 | 54.97 | 1.12 | 1.00 | 68.47 | 55.5 | 55.02 | 1.04 | 1.00 | 11.27 | 0.20 |
| fabp4 | 58.09 | 15 | 41.49 | 39.04 | 5.16 | 0.67 | 43.86 | 1 | 39.41 | 37.43 | 5.56 | 0.68 | 35.16 | 39.41 | 37.53 | 5.57 | 0.68 | 6.22 | 0.11 |
| fak1 | 54.7 | 7 | 36.8 | 35.92 | 4.02 | 0.66 | 86.09 | 6 | 26.85 | 25.54 | 3.64 | 0.49 | 60.03 | 23.87 | 22.67 | 3.65 | 0.44 | 12.93 | 0.24 |
| fgfr1 | 58.67 | 18 | 30.07 | 29.41 | 3.37 | 0.50 | 167.57 | 6 | 19.07 | 18.15 | 2.94 | 0.33 | 82.78 | 16.13 | 16.1 | 2.81 | 0.27 | 11 | 0.19 |
| fkb1a | 53.19 | 23 | 38.12 | 37.3 | 3.4 | 0.70 | 131.49 | 8 | 21.28 | 20.04 | 3.24 | 0.40 | 63.65 | 17.73 | 17.12 | 3.11 | 0.33 | 7.09 | 0.13 |
| fnta | 87.58 | 18 | 15.86 | 15.53 | 1.38 | 0.18 | 1528.55 | 3 | 7.93 | 7.8 | 1.05 | 0.09 | 583.78 | 7.09 | 7.21 | 1.05 | 0.08 | 4.05 | 0.05 |
| fpps | 97.77 | 14 | 81.47 | 80.23 | 3.73 | 0.82 | 184.69 | 2 | 72.7 | 72.4 | 4.73 | 0.74 | 86.18 | 66.43 | 66.5 | 4.56 | 0.68 | 1.25 | 0.01 |
| gcr | 69.67 | 20 | 27.09 | 26.77 | 2.68 | 0.38 | 324.83 | 1 | 17.42 | 17.65 | 2.39 | 0.25 | 138.15 | 17.42 | 17.6 | 2.52 | 0.25 | 7.74 | 0.11 |
| glcm | 70.24 | 19 | 61 | 58.32 | 4.15 | 0.83 | 67.52 | 3 | 36.97 | 36.23 | 5.27 | 0.53 | 44.52 | 35.12 | 33.2 | 5.63 | 0.50 | 16.64 | 0.24 |
| gria2 | 75.45 | 25 | 46.92 | 46.45 | 3.64 | 0.62 | 298.11 | 3 | 38.67 | 37.8 | 3.42 | 0.51 | 122.71 | 36.14 | 36.11 | 3.64 | 0.48 | 29.16 | 0.39 |
| grik1 | 65.78 | 13 | 56.81 | 56.38 | 2.7 | 0.86 | 146.64 | 1 | 30.9 | 31.25 | 4.18 | 0.47 | 71.32 | 30.9 | 31.64 | 4.25 | 0.47 | 16.94 | 0.26 |
| hdac2 | 57.73 | 21 | 42.19 | 41.52 | 2.81 | 0.72 | 240 | 5 | 29.42 | 29.04 | 2.6 | 0.51 | 103.94 | 28.87 | 28.68 | 2.82 | 0.50 | 21.09 | 0.37 |
| hdac8 | 63.06 | 12 | 63.06 | 62.26 | 1.28 | 0.99 | 208.54 | 6 | 45.21 | 43.92 | 3.01 | 0.72 | 111.70 | 32.72 | 32 | 2.88 | 0.52 | 36.88 | 0.58 |
| hivint | 67.98 | 18 | 16.99 | 16.36 | 3.46 | 0.24 | 151.65 | 3 | 9 | 8.64 | 2.82 | 0.13 | 70.73 | 6 | 6.03 | 2.36 | 0.09 | 1 | 0.01 |
| hivpr | 67.72 | 11 | 18.2 | 17.92 | 1.51 | 0.26 | 1043.03 | 4 | 12.76 | 12.67 | 1.37 | 0.19 | 396.82 | 17.26 | 17.09 | 1.46 | 0.25 | 0.75 | 0.01 |
| hivrt | 55.77 | 19 | 21.77 | 21.25 | 1.87 | 0.38 | 469.98 | 3 | 10.44 | 10.36 | 1.55 | 0.19 | 185.00 | 9.25 | 9.34 | 1.51 | 0.17 | 10.44 | 0.19 |
| hmdh | 52.33 | 16 | 46.45 | 45.27 | 2.32 | 0.87 | 208.51 | 2 | 31.16 | 30.52 | 2.64 | 0.60 | 93.81 | 25.87 | 25.18 | 2.66 | 0.49 | 4.12 | 0.08 |
| hs90a | 62.82 | 16 | 47.44 | 46.23 | 4.61 | 0.74 | 96.63 | 7 | 30.77 | 29.27 | 4.35 | 0.49 | 55.57 | 16.67 | 18.08 | 3.93 | 0.27 | 26.92 | 0.43 |
| hxk4 | 48.66 | 21 | 36.77 | 35.84 | 3.96 | 0.74 | 81.76 | 2 | 25.95 | 26.89 | 3.99 | 0.53 | 50.75 | 16.22 | 15.11 | 3.23 | 0.33 | 21.63 | 0.44 |
| igf1r | 63.61 | 25 | 32.14 | 30.98 | 3.11 | 0.49 | 236.86 | 3 | 19.42 | 19.24 | 2.94 | 0.31 | 94.26 | 18.75 | 17.89 | 2.75 | 0.29 | 10.71 | 0.17 |
| inha | 53.11 | 19 | 42.05 | 39.51 | 5.07 | 0.74 | 43.59 | 1 | 19.92 | 20.05 | 5.94 | 0.38 | 32.27 | 19.92 | 19.64 | 5.83 | 0.38 | 8.85 | 0.17 |
| ital | 62.3 | 13 | 24.63 | 24.33 | 3.42 | 0.39 | 166.16 | 2 | 21.01 | 20.41 | 3.23 | 0.34 | 93.50 | 15.21 | 15.33 | 2.84 | 0.24 | 3.62 | 0.06 |

| | | | | | | | | | | | | | | | | | | | |
|---|---|---|---|---|---|---|---|---|---|---|---|---|---|---|---|---|---|---|---|
| jak2 | 61.28 | 20 | 47.35 | 46.61 | 3.25 | 0.76 | 136.95 | 3 | 31.57 | 31.12 | 3.63 | 0.52 | 70.15 | 26 | 25.99 | 3.78 | 0.42 | 18.57 | 0.30 |
| kif11 | 59.9 | 19 | 51.34 | 50.32 | 3.23 | 0.84 | 145.57 | 4 | 48.77 | 47.95 | 2.84 | 0.81 | 71.95 | 44.49 | 44.84 | 3.29 | 0.74 | 3.42 | 0.06 |
| kit | 63.67 | 12 | 23.43 | 22.79 | 2.96 | 0.36 | 218.1 | 4 | 8.41 | 7.92 | 2.04 | 0.13 | 107.33 | 9.61 | 9.55 | 2.17 | 0.15 | 4.2 | 0.07 |
| kith | 50.91 | 34 | 50.91 | 50.35 | 1.53 | 0.99 | 64.06 | 1 | 50.91 | 49.49 | 2.59 | 1.00 | 36.73 | 50.91 | 49.39 | 2.75 | 1.00 | 28.85 | 0.57 |
| kpcb | 65.99 | 20 | 47.24 | 46.15 | 3.37 | 0.70 | 193.44 | 1 | 34.5 | 34.36 | 3.83 | 0.52 | 89.69 | 34.5 | 34.56 | 3.61 | 0.52 | 30.75 | 0.47 |
| lck | 66.11 | 16 | 36.99 | 36.64 | 1.86 | 0.55 | 789.45 | 2 | 19.81 | 19.69 | 1.81 | 0.30 | 298.70 | 17.66 | 17.81 | 1.66 | 0.27 | 16.94 | 0.26 |
| lkha4 | 45.6 | 16 | 28.43 | 27.43 | 2.42 | 0.60 | 164.68 | 3 | 21.32 | 20.97 | 2.48 | 0.47 | 82.32 | 17.77 | 18.45 | 2.57 | 0.39 | 10.07 | 0.22 |
| mapk2 | 62.12 | 24 | 61.13 | 59.57 | 2.23 | 0.96 | 141.52 | 4 | 49.3 | 49.04 | 3.32 | 0.79 | 66.43 | 41.41 | 40.83 | 3.96 | 0.67 | 35.5 | 0.57 |
| mcr | 82.51 | 19 | 34.54 | 34.13 | 6.01 | 0.41 | 71.77 | 2 | 15.35 | 15.19 | 4.87 | 0.19 | 48.13 | 15.35 | 15.57 | 5.09 | 0.19 | 9.59 | 0.12 |
| met | 68.51 | 9 | 43.27 | 42.91 | 3.25 | 0.63 | 249.09 | 4 | 30.65 | 29.97 | 3.13 | 0.45 | 115.02 | 34.86 | 35.23 | 3.56 | 0.51 | 7.81 | 0.11 |
| mk01 | 57.71 | 22 | 32.62 | 31.68 | 4.54 | 0.55 | 98.94 | 3 | 25.09 | 25.31 | 4.43 | 0.43 | 51.26 | 21.33 | 20.12 | 3.92 | 0.37 | 11.29 | 0.20 |
| mk10 | 64.21 | 19 | 36.42 | 35.25 | 4.05 | 0.55 | 145 | 4 | 23.96 | 23.28 | 3.62 | 0.37 | 69.18 | 17.25 | 17.22 | 3.37 | 0.27 | 14.38 | 0.22 |
| mk14 | 62.79 | 9 | 23.87 | 23.72 | 1.57 | 0.38 | 950.81 | 3 | 11.94 | 11.83 | 1.28 | 0.19 | 387.82 | 17.99 | 18.01 | 1.45 | 0.29 | 4.15 | 0.07 |
| mmp13 | 65.73 | 17 | 38.29 | 37.79 | 1.72 | 0.57 | 1176.44 | 1 | 18.01 | 18.07 | 1.47 | 0.27 | 438.33 | 39.86 | 39.68 | 1.81 | 0.61 | 6.99 | 0.11 |
| mp2k1 | 68.14 | 20 | 23.27 | 22.68 | 3.58 | 0.33 | 155.07 | 3 | 11.63 | 11.21 | 2.7 | 0.17 | 84.62 | 6.65 | 6.6 | 2.31 | 0.10 | 9.14 | 0.13 |
| nos1 | 81.82 | 20 | 44.44 | 43.39 | 4.51 | 0.53 | 159.34 | 4 | 30.3 | 29.54 | 4.28 | 0.37 | 85.36 | 34.34 | 34.09 | 4.47 | 0.42 | 24.24 | 0.30 |
| nram | 64.07 | 16 | 64.07 | 63.01 | 1.75 | 0.98 | 136.21 | 2 | 45.77 | 45.04 | 3.52 | 0.71 | 65.71 | 27.46 | 27.84 | 3.88 | 0.43 | 14.24 | 0.22 |
| pa2ga | 52.91 | 17 | 40.93 | 39.37 | 3.68 | 0.74 | 103.74 | 2 | 25.96 | 26.07 | 3.68 | 0.49 | 56.60 | 22.96 | 22.75 | 3.47 | 0.43 | 2 | 0.04 |
| parp1 | 60.03 | 26 | 58.26 | 57.94 | 0.8 | 0.97 | 971.19 | 2 | 45.86 | 45.64 | 1.36 | 0.76 | 339.21 | 40.54 | 40.51 | 1.56 | 0.68 | 39.95 | 0.67 |
| pde5a | 69.94 | 20 | 34.09 | 33.74 | 2.06 | 0.48 | 824.73 | 2 | 27.32 | 27.62 | 1.98 | 0.39 | 292.93 | 27.32 | 27.38 | 1.97 | 0.39 | 15.79 | 0.23 |
| pgh1 | 57.04 | 12 | 20.1 | 19.21 | 2.58 | 0.34 | 182.59 | 4 | 13.04 | 12.69 | 2.19 | 0.23 | 111.23 | 10.32 | 10.34 | 2.13 | 0.18 | 2.17 | 0.04 |
| pgh2 | 52.47 | 11 | 32.26 | 32.37 | 1.93 | 0.62 | 519.3 | 5 | 19.26 | 18.94 | 1.64 | 0.37 | 211.94 | 15.89 | 15.85 | 1.62 | 0.30 | 15.16 | 0.29 |
| plk1 | 64.84 | 23 | 42.29 | 40.71 | 3.62 | 0.63 | 152.91 | 3 | 35.71 | 35.38 | 3.85 | 0.55 | 74.55 | 27.25 | 26.8 | 3.97 | 0.42 | 17.85 | 0.28 |
| pnph | 68 | 27 | 65.13 | 63.32 | 2.74 | 0.93 | 142.01 | 2 | 51.72 | 51.23 | 4.12 | 0.76 | 72.89 | 48.85 | 47.96 | 4.13 | 0.72 | 45.01 | 0.66 |
| ppara | 48.87 | 9 | 19.6 | 19.6 | 1.76 | 0.40 | 428.92 | 3 | 11.55 | 11.63 | 1.51 | 0.24 | 190.85 | 6.44 | 6.38 | 1.2 | 0.13 | 3.76 | 0.08 |
| ppard | 50.17 | 6 | 23.22 | 23 | 2.25 | 0.46 | 241.61 | 1 | 12.44 | 12.78 | 1.98 | 0.25 | 121.94 | 12.44 | 12.79 | 1.97 | 0.25 | 0 | 0.00 |
| pparg | 53.06 | 20 | 22.39 | 22.13 | 1.61 | 0.42 | 777.48 | 1 | 10.78 | 10.74 | 1.21 | 0.20 | 249.12 | 10.78 | 10.71 | 1.29 | 0.20 | 3.32 | 0.06 |
| prgr | 58.68 | 25 | 45.33 | 44.64 | 2.39 | 0.76 | 318.85 | 5 | 21.45 | 20.93 | 2.33 | 0.37 | 144.63 | 18.21 | 18.25 | 2.19 | 0.31 | 29.95 | 0.51 |
| ptn1 | 56.37 | 14 | 38.85 | 38.06 | 3.54 | 0.68 | 146.85 | 2 | 29.71 | 29.87 | 3.13 | 0.53 | 78.34 | 26.66 | 26.94 | 3.4 | 0.47 | 7.62 | 0.14 |
| pur2 | 54.66 | 22 | 54.66 | 53.97 | 1.89 | 0.99 | 53.85 | 1 | 48.8 | 47.4 | 3.48 | 0.89 | 34.15 | 50.76 | 48.59 | 3.62 | 0.93 | 1.95 | 0.04 |
| pygm | 52 | 21 | 16.49 | 15.14 | 3.65 | 0.29 | 82.01 | 2 | 12.68 | 11.63 | 3.36 | 0.24 | 46.72 | 13.95 | 13.68 | 3.52 | 0.27 | 1.27 | 0.02 |
| pyrd | 55.87 | 14 | 50.55 | 49 | 3.21 | 0.88 | 109.65 | 5 | 31.93 | 30.59 | 3.53 | 0.57 | 69.18 | 31.04 | 31.07 | 4.34 | 0.56 | 10.64 | 0.19 |

| | | | | | | | | | | | | | | | | | | | |
|---|---|---|---|---|---|---|---|---|---|---|---|---|---|---|---|---|---|---|---|
| reni | 67.76 | 7 | 41.99 | 41.02 | 3.65 | 0.61 | 139.59 | 2 | 24.81 | 24.88 | 4.36 | 0.37 | 72.66 | 19.09 | 19.05 | 3.78 | 0.28 | 12.41 | 0.18 |
| rock1 | 63.74 | 26 | 40.83 | 39.75 | 4.13 | 0.62 | 122.19 | 5 | 25.89 | 24.96 | 3.53 | 0.41 | 68.16 | 15.94 | 15.95 | 3.18 | 0.25 | 19.92 | 0.31 |
| rxra | 53.25 | 21 | 53.25 | 52.87 | 0.89 | 0.99 | 127.75 | 4 | 39.11 | 38.61 | 3.06 | 0.73 | 69.61 | 32.45 | 32.95 | 3.09 | 0.61 | 34.11 | 0.64 |
| sahh | 48.02 | 28 | 48.02 | 47.98 | 0.33 | 1.00 | 56.47 | 2 | 48.02 | 46.06 | 2.83 | 1.00 | 36.47 | 46.36 | 45.74 | 2.78 | 0.97 | 38.08 | 0.79 |
| src | 66.49 | 12 | 32.67 | 32.22 | 1.68 | 0.48 | 955.4 | 2 | 17.96 | 18.02 | 1.48 | 0.27 | 399.24 | 14.52 | 14.85 | 1.44 | 0.22 | 9.17 | 0.14 |
| tgfr1 | 64.23 | 18 | 56.76 | 55.32 | 2.92 | 0.86 | 176.39 | 2 | 39.58 | 39.06 | 3.06 | 0.62 | 89.37 | 34.35 | 33.74 | 3.21 | 0.53 | 24.64 | 0.38 |
| thb | 66.72 | 24 | 53.38 | 51.4 | 3.76 | 0.77 | 135.8 | 2 | 42.09 | 41.42 | 4.41 | 0.63 | 68.22 | 32.85 | 33.81 | 4.05 | 0.49 | 22.58 | 0.34 |
| thrb | 59.91 | 7 | 45.92 | 45.66 | 1.61 | 0.76 | 678.75 | 3 | 32.36 | 32.47 | 1.75 | 0.54 | 270.53 | 25.36 | 25.56 | 1.7 | 0.42 | 7.22 | 0.12 |
| try1 | 58.56 | 9 | 58.34 | 58.05 | 0.59 | 0.99 | 693.85 | 2 | 53.44 | 53.41 | 1.12 | 0.91 | 262.63 | 50.54 | 50.56 | 1.62 | 0.86 | 6.01 | 0.10 |
| tryb1 | 52.41 | 15 | 43.68 | 42.65 | 2.38 | 0.81 | 155.78 | 1 | 25.53 | 25.06 | 2.9 | 0.49 | 81.97 | 25.53 | 25.08 | 2.93 | 0.49 | 7.39 | 0.14 |
| tysy | 62.61 | 20 | 35.39 | 34.42 | 3.93 | 0.55 | 156.22 | 4 | 25.41 | 24.3 | 3.41 | 0.41 | 74.01 | 19.96 | 20.03 | 3.58 | 0.32 | 14.52 | 0.23 |
| urok | 61.56 | 20 | 60.94 | 59.56 | 2.04 | 0.97 | 256.66 | 2 | 60.94 | 60.85 | 0.9 | 0.99 | 104.03 | 57.86 | 57.35 | 2.02 | 0.94 | 32.01 | 0.52 |
| vgfr2 | 57.28 | 10 | 26.78 | 26.6 | 1.85 | 0.46 | 509.54 | 2 | 15.87 | 15.93 | 1.71 | 0.28 | 244.17 | 15.62 | 15.52 | 1.68 | 0.27 | 5.95 | 0.10 |
| wee1 | 61.16 | 24 | 61.16 | 61.16 | 0 | 1.00 | 137.18 | 1 | 61.16 | 61.15 | 0.12 | 1.00 | 65.02 | 61.16 | 61.1 | 0.32 | 1.00 | 46.6 | 0.76 |
| xiap | 52.3 | 24 | 52.3 | 52.1 | 0.72 | 1.00 | 112.78 | 1 | 49.34 | 48.53 | 1.92 | 0.94 | 58.19 | 49.34 | 48.59 | 1.92 | 0.94 | 24.67 | 0.47 |
| Mean | 62.24 | 17.26 | 39.64 | 38.85 | 2.68 | 0.63 | 346.47 | 2.83 | 28.12 | 27.73 | 2.78 | 0.46 | 147.08 | 25.71 | 25.53 | 2.76 | 0.42 | 13.16 | 0.21 |

(a) Target identification on the DUD-E database (http://dude.docking.org/)

(b) EF at 1% of screened data for a perfect ranking of the active compounds on the ranked list

(c) Number of aggregated scoring functions

(d) EF of the best performing VS strategy

(e) Mean EF on 1000 bootstrap simulations of the best performing VS strategy

(f) Standard deviation of the bootstrap cross-validation procedure

(g) Fraction of the maximum possible EF that the VS method achieves

(h) Run time for each consensus scoring approach

Table TS4. VS performance of different scoring methods for the DUD-E database when BEDROC with α is set to 160.9 is used as model selection criterion and weighted scoring components are employed

| Target | CompScore | | | | | BISC | | | ASC |
|---|---|---|---|---|---|---|---|---|---|
| | Size | BEDROC | Boot. BEDROC | Std. Boot. BEDROC | Time | BEDROC | Boot. BEDROC | Std. Boot. BEDROC | BEDROC |
| aa2ar | 22 | 0.74 | 0.74 | 0.02 | 1388.29 | 0.55 | 0.55 | 0.02 | 0.35 |
| abl1 | 19 | 0.62 | 0.62 | 0.04 | 370.36 | 0.49 | 0.48 | 0.04 | 0.19 |
| ace | 21 | 0.54 | 0.54 | 0.03 | 799.7 | 0.3 | 0.3 | 0.03 | 0.04 |
| aces | 14 | 0.71 | 0.71 | 0.02 | 916.06 | 0.46 | 0.46 | 0.03 | 0.15 |
| ada | 17 | 0.77 | 0.77 | 0.04 | 135.22 | 0.33 | 0.33 | 0.06 | 0.07 |
| ada17 | 14 | 0.79 | 0.79 | 0.01 | 1832.18 | 0.7 | 0.7 | 0.02 | 0.37 |
| adrb1 | 14 | 0.53 | 0.53 | 0.04 | 589.61 | 0.16 | 0.16 | 0.03 | 0.13 |
| adrb2 | 19 | 0.44 | 0.43 | 0.04 | 594.32 | 0.23 | 0.23 | 0.04 | 0.15 |
| akt1 | 24 | 0.71 | 0.7 | 0.03 | 812.16 | 0.45 | 0.45 | 0.03 | 0.24 |
| akt2 | 21 | 0.7 | 0.7 | 0.04 | 184.32 | 0.43 | 0.42 | 0.06 | 0.11 |
| aldr | 23 | 0.68 | 0.68 | 0.04 | 371.02 | 0.54 | 0.54 | 0.04 | 0.04 |
| ampc | 15 | 0.72 | 0.72 | 0.07 | 69.25 | 0.16 | 0.16 | 0.07 | 0.01 |
| andr | 15 | 0.6 | 0.6 | 0.03 | 420.84 | 0.28 | 0.28 | 0.04 | 0.17 |
| aofb | 18 | 0.42 | 0.42 | 0.06 | 164.75 | 0.21 | 0.21 | 0.05 | 0.06 |
| bace1 | 28 | 0.33 | 0.32 | 0.04 | 870.06 | 0.18 | 0.18 | 0.03 | 0.13 |
| braf | 9 | 0.61 | 0.61 | 0.04 | 302.52 | 0.35 | 0.35 | 0.04 | 0.21 |
| cah2 | 25 | 0.54 | 0.54 | 0.02 | 1500.49 | 0.17 | 0.17 | 0.02 | 0.05 |
| casp3 | 17 | 0.41 | 0.4 | 0.05 | 392.34 | 0.19 | 0.19 | 0.04 | 0.04 |
| cdk2 | 21 | 0.51 | 0.51 | 0.02 | 1070.25 | 0.34 | 0.33 | 0.03 | 0.21 |
| comt | 26 | 0.99 | 0.99 | 0.01 | 98.61 | 0.92 | 0.92 | 0.03 | 0.15 |
| cp2c9 | 31 | 0.26 | 0.26 | 0.05 | 250.77 | 0.11 | 0.11 | 0.04 | 0.06 |
| cp3a4 | 22 | 0.25 | 0.25 | 0.04 | 466.11 | 0.16 | 0.16 | 0.04 | 0.06 |
| csf1r | 13 | 0.59 | 0.59 | 0.04 | 353.53 | 0.35 | 0.35 | 0.04 | 0.11 |

| | | | | | | | | | |
|---|---|---|---|---|---|---|---|---|---|
| cxcr4 | 27 | 0.73 | 0.72 | 0.06 | 87.07 | 0.3 | 0.29 | 0.09 | 0.12 |
| def | 18 | 0.91 | 0.91 | 0.02 | 168.79 | 0.74 | 0.74 | 0.04 | 0.1 |
| dhi1 | 17 | 0.42 | 0.41 | 0.03 | 656.72 | 0.19 | 0.19 | 0.03 | 0.07 |
| dpp4 | 27 | 0.66 | 0.66 | 0.02 | 2385.56 | 0.5 | 0.5 | 0.02 | 0.17 |
| drd3 | 21 | 0.45 | 0.45 | 0.03 | 1728.59 | 0.17 | 0.17 | 0.02 | 0.02 |
| dyr | 22 | 0.76 | 0.76 | 0.03 | 752.26 | 0.61 | 0.61 | 0.03 | 0.36 |
| egfr | 13 | 0.75 | 0.75 | 0.02 | 1696.56 | 0.43 | 0.43 | 0.02 | 0.34 |
| esr1 | 19 | 0.79 | 0.78 | 0.02 | 931.6 | 0.67 | 0.66 | 0.03 | 0.09 |
| esr2 | 20 | 0.8 | 0.79 | 0.02 | 892.88 | 0.6 | 0.6 | 0.03 | 0.12 |
| fa10 | 15 | 0.75 | 0.75 | 0.02 | 1404.37 | 0.59 | 0.59 | 0.02 | 0.2 |
| fa7 | 17 | 0.95 | 0.95 | 0.01 | 167.68 | 0.9 | 0.9 | 0.02 | 0.23 |
| fabp4 | 24 | 0.87 | 0.87 | 0.04 | 46.35 | 0.7 | 0.69 | 0.07 | 0.15 |
| fak1 | 14 | 0.71 | 0.71 | 0.05 | 133.74 | 0.48 | 0.48 | 0.06 | 0.3 |
| fgfr1 | 24 | 0.57 | 0.57 | 0.05 | 260.84 | 0.26 | 0.26 | 0.05 | 0.25 |
| fkb1a | 26 | 0.73 | 0.73 | 0.04 | 182.83 | 0.34 | 0.34 | 0.06 | 0.15 |
| fnta | 21 | 0.27 | 0.27 | 0.02 | 3158.1 | 0.1 | 0.1 | 0.01 | 0.04 |
| fpps | 16 | 0.94 | 0.93 | 0.02 | 284.27 | 0.87 | 0.87 | 0.03 | 0.02 |
| gcr | 17 | 0.54 | 0.54 | 0.04 | 445.71 | 0.29 | 0.29 | 0.04 | 0.12 |
| glcm | 24 | 0.86 | 0.85 | 0.04 | 94.21 | 0.39 | 0.39 | 0.07 | 0.09 |
| gria2 | 31 | 0.73 | 0.72 | 0.03 | 499.57 | 0.57 | 0.57 | 0.04 | 0.43 |
| grik1 | 20 | 0.82 | 0.82 | 0.03 | 313.45 | 0.55 | 0.55 | 0.06 | 0.32 |
| hdac2 | 22 | 0.75 | 0.75 | 0.03 | 441.63 | 0.44 | 0.44 | 0.04 | 0.33 |
| hdac8 | 16 | 0.9 | 0.9 | 0.02 | 343.87 | 0.52 | 0.52 | 0.04 | 0.48 |
| hivint | 19 | 0.4 | 0.4 | 0.05 | 178.8 | 0.14 | 0.14 | 0.04 | 0.03 |
| hivpr | 6 | 0.45 | 0.45 | 0.02 | 1395.58 | 0.29 | 0.29 | 0.02 | 0.02 |
| hivrt | 15 | 0.46 | 0.46 | 0.03 | 807.05 | 0.18 | 0.18 | 0.03 | 0.19 |
| hmdh | 19 | 0.86 | 0.86 | 0.02 | 276.17 | 0.4 | 0.4 | 0.05 | 0.11 |
| hs90a | 21 | 0.76 | 0.75 | 0.05 | 108.64 | 0.2 | 0.2 | 0.05 | 0.27 |
| hxk4 | 18 | 0.78 | 0.78 | 0.04 | 94.63 | 0.48 | 0.47 | 0.07 | 0.45 |
| igf1r | 24 | 0.59 | 0.59 | 0.04 | 253.57 | 0.34 | 0.34 | 0.05 | 0.24 |
| inha | 21 | 0.72 | 0.71 | 0.07 | 47.37 | 0.45 | 0.44 | 0.1 | 0.19 |

| | | | | | | | | | |
|---|---|---|---|---|---|---|---|---|---|
| ital | 25 | 0.47 | 0.47 | 0.05 | 226.85 | 0.28 | 0.28 | 0.05 | 0.09 |
| jak2 | 26 | 0.85 | 0.84 | 0.03 | 167.85 | 0.51 | 0.51 | 0.06 | 0.35 |
| kif11 | 27 | 0.87 | 0.87 | 0.02 | 181.51 | 0.77 | 0.77 | 0.04 | 0.16 |
| kit | 18 | 0.41 | 0.41 | 0.05 | 365.22 | 0.16 | 0.16 | 0.03 | 0.07 |
| kith | 32 | 0.97 | 0.97 | 0.01 | 72.85 | 0.88 | 0.88 | 0.03 | 0.62 |
| kpcb | 18 | 0.77 | 0.77 | 0.03 | 271.21 | 0.62 | 0.62 | 0.05 | 0.53 |
| lck | 15 | 0.63 | 0.63 | 0.02 | 1074.97 | 0.31 | 0.31 | 0.03 | 0.29 |
| lkha4 | 10 | 0.66 | 0.66 | 0.04 | 171.64 | 0.43 | 0.43 | 0.05 | 0.22 |
| mapk2 | 35 | 0.93 | 0.93 | 0.02 | 169.57 | 0.7 | 0.7 | 0.05 | 0.66 |
| mcr | 11 | 0.65 | 0.65 | 0.06 | 70.06 | 0.25 | 0.24 | 0.08 | 0.1 |
| met | 20 | 0.71 | 0.71 | 0.03 | 315.02 | 0.61 | 0.61 | 0.04 | 0.15 |
| mk01 | 21 | 0.64 | 0.64 | 0.06 | 120.41 | 0.34 | 0.34 | 0.06 | 0.23 |
| mk10 | 19 | 0.57 | 0.57 | 0.05 | 148.8 | 0.29 | 0.29 | 0.06 | 0.23 |
| mk14 | 20 | 0.45 | 0.45 | 0.03 | 1800.61 | 0.35 | 0.35 | 0.03 | 0.07 |
| mmp13 | 18 | 0.74 | 0.74 | 0.02 | 1957.67 | 0.56 | 0.56 | 0.02 | 0.17 |
| mp2k1 | 23 | 0.44 | 0.44 | 0.05 | 278.87 | 0.09 | 0.09 | 0.04 | 0.13 |
| nos1 | 25 | 0.69 | 0.69 | 0.04 | 262.55 | 0.38 | 0.38 | 0.06 | 0.38 |
| nram | 24 | 0.95 | 0.95 | 0.01 | 178.38 | 0.47 | 0.47 | 0.05 | 0.34 |
| pa2ga | 22 | 0.76 | 0.75 | 0.04 | 119.99 | 0.45 | 0.45 | 0.06 | 0.06 |
| parp1 | 26 | 0.91 | 0.9 | 0.01 | 1388.88 | 0.71 | 0.71 | 0.02 | 0.62 |
| pde5a | 22 | 0.59 | 0.59 | 0.03 | 1351.57 | 0.47 | 0.47 | 0.03 | 0.31 |
| pgh1 | 13 | 0.41 | 0.4 | 0.05 | 356.61 | 0.2 | 0.2 | 0.04 | 0.05 |
| pgh2 | 12 | 0.65 | 0.65 | 0.03 | 834.71 | 0.33 | 0.33 | 0.03 | 0.25 |
| plk1 | 27 | 0.74 | 0.74 | 0.04 | 329.42 | 0.49 | 0.49 | 0.05 | 0.31 |
| pnph | 26 | 0.92 | 0.92 | 0.02 | 168.52 | 0.63 | 0.63 | 0.05 | 0.64 |
| ppara | 13 | 0.42 | 0.42 | 0.03 | 509 | 0.09 | 0.09 | 0.02 | 0.08 |
| ppard | 10 | 0.53 | 0.53 | 0.04 | 356.9 | 0.16 | 0.16 | 0.03 | 0 |
| pparg | 21 | 0.49 | 0.49 | 0.03 | 1016.67 | 0.22 | 0.22 | 0.03 | 0.05 |
| prgr | 25 | 0.77 | 0.77 | 0.02 | 571.18 | 0.34 | 0.34 | 0.04 | 0.4 |
| ptn1 | 16 | 0.72 | 0.72 | 0.04 | 231.03 | 0.49 | 0.49 | 0.05 | 0.17 |
| pur2 | 21 | 0.97 | 0.97 | 0.01 | 51.42 | 0.74 | 0.73 | 0.06 | 0.04 |

| | | | | | | | | | |
|---|---|---|---|---|---|---|---|---|---|
| pygm | 19 | 0.51 | 0.51 | 0.07 | 83.16 | 0.22 | 0.23 | 0.06 | 0 |
| pyrd | 19 | 0.84 | 0.84 | 0.03 | 143.15 | 0.61 | 0.61 | 0.05 | 0.23 |
| reni | 11 | 0.66 | 0.66 | 0.05 | 160.31 | 0.38 | 0.37 | 0.06 | 0.24 |
| rock1 | 20 | 0.72 | 0.71 | 0.04 | 174.39 | 0.29 | 0.29 | 0.05 | 0.29 |
| rxra | 21 | 0.95 | 0.95 | 0.01 | 131.3 | 0.65 | 0.65 | 0.05 | 0.69 |
| sahh | 22 | 0.98 | 0.98 | 0.01 | 45.68 | 0.91 | 0.9 | 0.03 | 0.54 |
| src | 13 | 0.55 | 0.56 | 0.02 | 1596.35 | 0.26 | 0.26 | 0.02 | 0.15 |
| tgfr1 | 18 | 0.87 | 0.87 | 0.02 | 272.91 | 0.58 | 0.57 | 0.04 | 0.43 |
| thb | 14 | 0.83 | 0.83 | 0.03 | 132.72 | 0.56 | 0.55 | 0.05 | 0.45 |
| thrb | 15 | 0.77 | 0.77 | 0.02 | 1122.75 | 0.44 | 0.44 | 0.03 | 0.11 |
| try1 | 16 | 0.92 | 0.92 | 0.01 | 1158.06 | 0.77 | 0.77 | 0.02 | 0.12 |
| tryb1 | 16 | 0.81 | 0.81 | 0.03 | 241.66 | 0.54 | 0.54 | 0.05 | 0.11 |
| tysy | 20 | 0.65 | 0.65 | 0.05 | 185.15 | 0.38 | 0.38 | 0.06 | 0.26 |
| urok | 20 | 0.94 | 0.94 | 0.01 | 439.94 | 0.89 | 0.89 | 0.02 | 0.44 |
| vgfr2 | 14 | 0.53 | 0.53 | 0.03 | 958.08 | 0.33 | 0.33 | 0.03 | 0.14 |
| wee1 | 22 | 1 | 1 | 0 | 156.2 | 0.97 | 0.97 | 0.01 | 0.7 |
| xiap | 24 | 0.97 | 0.97 | 0.01 | 124.81 | 0.87 | 0.87 | 0.03 | 0.56 |
| Mean | 19.68 | 0.69 | 0.68 | 0.03 | 553.31 | 0.44 | 0.44 | 0.04 | 0.22 |

Table TS5. VS performance of different scoring methods for the DUD-E database when EF at the first 1% of screened data is used as model selection criterion and weighted scoring components are employed

| Target[a] | Opt. EF[b] | CompScore | | | | | | BISC | | | | ASC | |
|---|---|---|---|---|---|---|---|---|---|---|---|---|---|
| | | Size[c] | EF[d] | Boot. EF[e] | Std. Boot. EF[f] | Fract. Opt. EF[g] | Time (seconds) [h] | EF[d] | Boot. EF[e] | Std. Boot. EF[f] | Fract. Opt. EF[g] | EF[d] | Fract. Opt. EF[g] |
| aa2ar | 66.17 | 27 | 46.61 | 46.3 | 1.69 | 0.70 | 1360.07 | 33.71 | 33.97 | 1.71 | 0.51 | 19.35 | 0.29 |
| abl1 | 59.61 | 24 | 29.81 | 28.81 | 2.84 | 0.50 | 343.29 | 25.39 | 24.98 | 2.63 | 0.43 | 11.59 | 0.19 |
| ace | 60.77 | 24 | 29.14 | 28.71 | 2.32 | 0.48 | 707.33 | 15.99 | 16.16 | 2.05 | 0.26 | 1.78 | 0.03 |
| aces | 51.64 | 15 | 35.86 | 35.65 | 1.92 | 0.69 | 835.67 | 20.52 | 20.23 | 1.65 | 0.40 | 6.99 | 0.14 |
| ada | 59.35 | 31 | 44.52 | 42.78 | 3.63 | 0.75 | 176.7 | 19.08 | 18.68 | 3.43 | 0.32 | 3.18 | 0.05 |
| ada17 | 68.15 | 26 | 51.25 | 51.04 | 1.68 | 0.75 | 1621.2 | 42.8 | 42.83 | 1.97 | 0.63 | 21.59 | 0.32 |
| adrb1 | 62.23 | 32 | 27.88 | 27.66 | 2.38 | 0.45 | 664.69 | 9.29 | 9.23 | 1.79 | 0.15 | 8.49 | 0.14 |
| adrb2 | 58.77 | 23 | 24.82 | 24.03 | 2.43 | 0.42 | 511.15 | 11.75 | 11.64 | 2.05 | 0.20 | 6.97 | 0.12 |
| akt1 | 56.53 | 22 | 38.61 | 38.11 | 2.23 | 0.68 | 640.26 | 23.44 | 23.34 | 2.05 | 0.41 | 11.72 | 0.21 |
| akt2 | 59.36 | 24 | 39.57 | 39.04 | 4.14 | 0.67 | 236.88 | 23.23 | 22.8 | 3.69 | 0.39 | 6.88 | 0.12 |
| aldr | 57.13 | 20 | 38.93 | 38.09 | 2.93 | 0.68 | 345 | 27 | 27.69 | 2.95 | 0.47 | 2.51 | 0.04 |
| ampc | 59.94 | 25 | 37.2 | 34.5 | 5.37 | 0.62 | 78.06 | 8.27 | 7.57 | 3.54 | 0.14 | 0 | 0.00 |
| andr | 59.84 | 22 | 34.19 | 33.47 | 2.54 | 0.57 | 432.27 | 13.05 | 13.16 | 2.24 | 0.22 | 7.65 | 0.13 |
| aofb | 57.55 | 36 | 16.68 | 15.94 | 3.07 | 0.29 | 212.74 | 11.68 | 10.75 | 2.65 | 0.20 | 3.34 | 0.06 |
| bace1 | 65.61 | 26 | 17.11 | 17.02 | 2.06 | 0.26 | 755.62 | 9.63 | 9.45 | 1.79 | 0.15 | 7.13 | 0.11 |
| braf | 66.15 | 28 | 32.09 | 31.09 | 3.07 | 0.49 | 388.47 | 19.65 | 19.56 | 2.85 | 0.30 | 11.79 | 0.18 |
| cah2 | 64.1 | 28 | 32.35 | 32.36 | 1.73 | 0.50 | 1539.16 | 9.97 | 9.79 | 1.27 | 0.16 | 3.05 | 0.05 |
| casp3 | 54.64 | 31 | 14.54 | 14.14 | 2.23 | 0.27 | 424.39 | 10.03 | 9.82 | 1.93 | 0.18 | 3.01 | 0.06 |
| cdk2 | 59.56 | 26 | 29.25 | 28.93 | 1.73 | 0.49 | 1126.07 | 17.05 | 16.91 | 1.49 | 0.29 | 12.63 | 0.21 |
| comt | 94.27 | 30 | 91.85 | 90.53 | 3.1 | 0.97 | 120.68 | 79.77 | 78.29 | 4.9 | 0.85 | 7.25 | 0.08 |
| cp2c9 | 61.89 | 40 | 15.05 | 13.84 | 3.07 | 0.24 | 300.88 | 5.85 | 5.41 | 1.95 | 0.09 | 4.18 | 0.07 |

| | | | | | | | | | | | | | |
|---|---|---|---|---|---|---|---|---|---|---|---|---|---|
| cp3a4 | 70.18 | 32 | 12.87 | 12.58 | 2.46 | 0.18 | 476.93 | 10.53 | 10.21 | 2.19 | 0.15 | 2.92 | 0.04 |
| csf1r | 73.81 | 16 | 36.61 | 35.74 | 3.04 | 0.50 | 411.72 | 24 | 23.7 | 2.85 | 0.33 | 6.6 | 0.09 |
| cxcr4 | 86.02 | 27 | 51.61 | 49.64 | 6.45 | 0.60 | 100.45 | 19.66 | 19.46 | 6.01 | 0.23 | 9.83 | 0.11 |
| def | 58.46 | 21 | 58.46 | 57.01 | 2.39 | 1.00 | 182.59 | 37.3 | 37.85 | 3.59 | 0.64 | 5.04 | 0.09 |
| dhi1 | 58.9 | 26 | 19.63 | 19.46 | 2.04 | 0.33 | 749.89 | 8.76 | 8.82 | 1.5 | 0.15 | 3.62 | 0.06 |
| dpp4 | 77.65 | 33 | 44.45 | 43.87 | 1.73 | 0.57 | 2007.78 | 33.38 | 33.42 | 1.84 | 0.43 | 10.5 | 0.14 |
| drd3 | 70.5 | 27 | 27.24 | 26.89 | 1.78 | 0.39 | 1442.11 | 9.77 | 10.11 | 1.32 | 0.14 | 1.66 | 0.02 |
| dyr | 75.66 | 25 | 52.18 | 51.52 | 2.75 | 0.69 | 749.7 | 39.14 | 39.24 | 2.65 | 0.52 | 22.18 | 0.29 |
| egfr | 65.57 | 13 | 48.95 | 48.67 | 1.52 | 0.75 | 1431.16 | 25.86 | 25.4 | 1.57 | 0.39 | 19.39 | 0.30 |
| esr1 | 54.86 | 23 | 46.16 | 45.57 | 1.81 | 0.84 | 786.89 | 34.29 | 34.05 | 2.07 | 0.63 | 4.48 | 0.08 |
| esr2 | 55.88 | 20 | 46.34 | 45.76 | 1.64 | 0.83 | 773.7 | 29.99 | 29.95 | 1.96 | 0.54 | 6.54 | 0.12 |
| fa10 | 53.51 | 22 | 41.2 | 40.76 | 1.5 | 0.77 | 1213.58 | 28.71 | 28.59 | 1.48 | 0.54 | 10.63 | 0.20 |
| fa7 | 55.5 | 28 | 55.5 | 55.09 | 1.06 | 1.00 | 211.04 | 55.5 | 55 | 1.12 | 1.00 | 12.14 | 0.22 |
| fabp4 | 58.09 | 29 | 47.71 | 46.56 | 5.71 | 0.82 | 71.02 | 39.41 | 37.53 | 5.73 | 0.68 | 12.45 | 0.21 |
| fak1 | 54.7 | 23 | 32.82 | 32.59 | 4.25 | 0.60 | 162.36 | 23.87 | 22.58 | 3.45 | 0.44 | 14.92 | 0.27 |
| fgfr1 | 58.67 | 33 | 30.8 | 29.52 | 3.3 | 0.52 | 275.61 | 16.13 | 16.18 | 2.87 | 0.27 | 11.73 | 0.20 |
| fkb1a | 53.19 | 26 | 41.66 | 40.14 | 3.36 | 0.78 | 193.73 | 17.73 | 17.45 | 3.14 | 0.33 | 7.09 | 0.13 |
| fnta | 87.58 | 29 | 15.69 | 15.44 | 1.37 | 0.18 | 2605.19 | 7.09 | 7.17 | 1.04 | 0.08 | 3.04 | 0.03 |
| fpps | 97.77 | 18 | 86.49 | 86.15 | 3.46 | 0.88 | 320.17 | 76.46 | 76.25 | 4.39 | 0.78 | 1.25 | 0.01 |
| gcr | 69.67 | 23 | 30.48 | 30.1 | 2.81 | 0.44 | 490.28 | 17.42 | 17.54 | 2.33 | 0.25 | 7.74 | 0.11 |
| glcm | 70.24 | 30 | 61 | 58.69 | 4.87 | 0.87 | 114.7 | 25.88 | 23.56 | 4.97 | 0.37 | 5.55 | 0.08 |
| gria2 | 75.45 | 32 | 48.82 | 48.78 | 3.83 | 0.65 | 520.88 | 36.14 | 35.81 | 3.43 | 0.48 | 25.36 | 0.34 |
| grik1 | 65.78 | 38 | 43.85 | 42.43 | 4.01 | 0.67 | 278.72 | 30.9 | 31.63 | 4.32 | 0.47 | 16.94 | 0.26 |
| hdac2 | 57.73 | 31 | 41.08 | 40.44 | 3.11 | 0.71 | 428.48 | 28.31 | 28.29 | 2.88 | 0.49 | 18.87 | 0.33 |
| hdac8 | 63.06 | 19 | 63.06 | 61.95 | 1.55 | 1.00 | 401.33 | 32.72 | 32.2 | 2.84 | 0.52 | 26.77 | 0.42 |
| hivint | 67.98 | 32 | 20.99 | 19.55 | 3.57 | 0.31 | 281.91 | 9 | 8.84 | 2.68 | 0.13 | 3 | 0.04 |
| hivpr | 67.72 | 14 | 24.95 | 24.6 | 1.61 | 0.37 | 1561.48 | 17.26 | 17.13 | 1.42 | 0.25 | 0.94 | 0.01 |
| hivrt | 55.77 | 26 | 22.37 | 22.09 | 1.96 | 0.40 | 696.29 | 8.65 | 8.69 | 1.43 | 0.16 | 9.25 | 0.17 |
| hmdh | 52.33 | 21 | 48.8 | 47.84 | 1.89 | 0.93 | 313.26 | 19.4 | 19.41 | 2.56 | 0.37 | 5.29 | 0.10 |
| hs90a | 62.82 | 22 | 46.15 | 45.72 | 4.97 | 0.73 | 142.66 | 14.1 | 14.91 | 3.68 | 0.22 | 19.23 | 0.31 |
| hxk4 | 48.66 | 21 | 40.01 | 39.05 | 3.88 | 0.82 | 117.68 | 20.55 | 20.62 | 4.29 | 0.42 | 20.55 | 0.42 |

| | | | | | | | | | | | | | |
|---|---|---|---|---|---|---|---|---|---|---|---|---|---|
| igf1r | 63.61 | 30 | 34.82 | 33.83 | 3.13 | 0.55 | 353.42 | 18.75 | 17.88 | 2.73 | 0.29 | 11.38 | 0.18 |
| inha | 53.11 | 32 | 39.84 | 37.86 | 5.68 | 0.75 | 59.64 | 19.92 | 20.01 | 5.69 | 0.38 | 8.85 | 0.17 |
| ital | 62.3 | 33 | 23.9 | 22.84 | 3.03 | 0.38 | 314.91 | 15.21 | 15.39 | 2.88 | 0.24 | 5.07 | 0.08 |
| jak2 | 61.28 | 25 | 52.92 | 52.02 | 3.3 | 0.86 | 226.61 | 26 | 25.68 | 3.74 | 0.42 | 19.5 | 0.32 |
| kif11 | 59.9 | 27 | 55.62 | 53.86 | 2.95 | 0.93 | 254.08 | 44.49 | 44.89 | 3.25 | 0.74 | 10.27 | 0.17 |
| kit | 63.67 | 24 | 19.22 | 18.42 | 2.65 | 0.30 | 389.06 | 9.61 | 9.38 | 2.2 | 0.15 | 3.6 | 0.06 |
| kith | 50.91 | 39 | 50.91 | 50.54 | 1.12 | 1.00 | 95.72 | 50.91 | 49.42 | 2.73 | 1.00 | 28.85 | 0.57 |
| kpcb | 65.99 | 33 | 47.24 | 46.79 | 3.7 | 0.72 | 330.47 | 34.5 | 34.56 | 3.78 | 0.52 | 30.75 | 0.47 |
| lck | 66.11 | 21 | 36.75 | 36.5 | 1.96 | 0.56 | 1175.48 | 17.66 | 17.8 | 1.67 | 0.27 | 16.23 | 0.25 |
| lkha4 | 45.6 | 27 | 26.06 | 25.46 | 2.73 | 0.57 | 250.72 | 17.77 | 18.41 | 2.51 | 0.39 | 11.25 | 0.25 |
| mapk2 | 62.12 | 39 | 61.13 | 60.1 | 1.78 | 0.98 | 244.36 | 41.41 | 40.73 | 3.8 | 0.67 | 38.46 | 0.62 |
| mcr | 82.51 | 19 | 38.38 | 37.19 | 5.84 | 0.47 | 95.18 | 15.35 | 15.14 | 4.97 | 0.19 | 5.76 | 0.07 |
| met | 68.51 | 22 | 45.67 | 44.66 | 3.02 | 0.67 | 387.48 | 34.86 | 35.17 | 3.45 | 0.51 | 9.62 | 0.14 |
| mk01 | 57.71 | 45 | 32.62 | 30.95 | 4.5 | 0.57 | 165.74 | 21.33 | 20.24 | 4.02 | 0.37 | 11.29 | 0.20 |
| mk10 | 64.21 | 27 | 37.38 | 35.84 | 3.95 | 0.58 | 250.5 | 16.29 | 16.33 | 3.24 | 0.25 | 12.46 | 0.19 |
| mk14 | 62.79 | 21 | 25.08 | 24.97 | 1.56 | 0.40 | 1519.57 | 17.99 | 18.07 | 1.43 | 0.29 | 3.63 | 0.06 |
| mmp13 | 65.73 | 21 | 46.85 | 46.78 | 1.61 | 0.71 | 1684.44 | 34.79 | 34.64 | 1.61 | 0.53 | 10.84 | 0.16 |
| mp2k1 | 68.14 | 21 | 24.93 | 24.06 | 3.66 | 0.37 | 276.09 | 6.65 | 6.57 | 2.27 | 0.10 | 9.14 | 0.13 |
| nos1 | 81.82 | 32 | 47.47 | 45.67 | 4.1 | 0.58 | 319.46 | 24.24 | 24.26 | 3.76 | 0.30 | 22.22 | 0.27 |
| nram | 64.07 | 35 | 63.05 | 62.16 | 1.85 | 0.98 | 246.09 | 27.46 | 27.65 | 3.72 | 0.43 | 17.29 | 0.27 |
| pa2ga | 52.91 | 23 | 41.93 | 40.68 | 3.93 | 0.79 | 153 | 22.96 | 22.76 | 3.35 | 0.43 | 2.99 | 0.06 |
| parp1 | 60.03 | 44 | 57.67 | 57.16 | 1.12 | 0.96 | 1542.55 | 40.54 | 40.46 | 1.61 | 0.68 | 32.87 | 0.55 |
| pde5a | 69.94 | 35 | 35.34 | 34.94 | 1.87 | 0.51 | 1241.17 | 27.32 | 27.28 | 1.95 | 0.39 | 17.8 | 0.25 |
| pgh1 | 57.04 | 17 | 20.1 | 19.87 | 2.87 | 0.35 | 310.05 | 9.24 | 9.5 | 2.02 | 0.16 | 2.17 | 0.04 |
| pgh2 | 52.47 | 22 | 32.5 | 32.13 | 1.8 | 0.62 | 808.63 | 15.89 | 15.84 | 1.59 | 0.30 | 13 | 0.25 |
| plk1 | 64.84 | 30 | 46.99 | 45.36 | 3.3 | 0.72 | 229.59 | 27.25 | 26.82 | 4 | 0.42 | 16.91 | 0.26 |
| pnph | 68 | 30 | 66.08 | 64.68 | 2.61 | 0.97 | 235.27 | 40.23 | 40.17 | 4.26 | 0.59 | 36.39 | 0.54 |
| ppara | 48.87 | 22 | 19.06 | 18.7 | 1.75 | 0.39 | 602.02 | 3.76 | 3.68 | 0.98 | 0.08 | 3.49 | 0.07 |
| ppard | 50.17 | 10 | 24.88 | 24.55 | 2.23 | 0.50 | 371.88 | 8.29 | 8.25 | 1.7 | 0.17 | 0 | 0.00 |
| pparg | 53.06 | 28 | 24.67 | 24.31 | 1.64 | 0.46 | 1074.09 | 10.78 | 10.77 | 1.31 | 0.20 | 2.9 | 0.05 |
| prgr | 58.68 | 24 | 46.95 | 45.93 | 2.01 | 0.80 | 536.39 | 17.81 | 18.23 | 2.1 | 0.30 | 19.02 | 0.32 |

| | | | | | | | | | | | | | |
|---|---|---|---|---|---|---|---|---|---|---|---|---|---|
| ptn1 | 56.37 | 18 | 39.61 | 39.07 | 3.87 | 0.70 | 220.75 | 26.66 | 26.92 | 3.4 | 0.47 | 7.62 | 0.14 |
| pur2 | 54.66 | 34 | 54.66 | 53.47 | 2.65 | 1.00 | 70.3 | 39.04 | 38.83 | 4.74 | 0.71 | 1.95 | 0.04 |
| pygm | 52 | 29 | 16.49 | 15.1 | 3.75 | 0.32 | 118.73 | 13.95 | 13.8 | 3.46 | 0.27 | 0 | 0.00 |
| pyrd | 55.87 | 27 | 50.55 | 49.15 | 3.38 | 0.90 | 179.65 | 31.04 | 31.1 | 4.26 | 0.56 | 10.64 | 0.19 |
| reni | 67.76 | 17 | 37.22 | 36.64 | 4.04 | 0.55 | 201.44 | 19.09 | 18.97 | 3.88 | 0.28 | 14.32 | 0.21 |
| rock1 | 63.74 | 26 | 43.82 | 41.46 | 3.86 | 0.69 | 212.65 | 15.94 | 15.94 | 3.31 | 0.25 | 16.93 | 0.27 |
| rxra | 53.25 | 25 | 53.25 | 52.9 | 0.94 | 1.00 | 166.48 | 32.45 | 33 | 3.17 | 0.61 | 35.78 | 0.67 |
| sahh | 48.02 | 34 | 48.02 | 47.65 | 1.03 | 1.00 | 70 | 46.36 | 45.59 | 2.87 | 0.97 | 28.15 | 0.59 |
| src | 66.49 | 14 | 34.01 | 33.95 | 1.71 | 0.51 | 1373.12 | 14.52 | 14.66 | 1.43 | 0.22 | 8.22 | 0.12 |
| tgfr1 | 64.23 | 29 | 59 | 57.11 | 2.68 | 0.92 | 288.13 | 34.35 | 33.81 | 3.1 | 0.53 | 23.15 | 0.36 |
| thb | 66.72 | 32 | 55.43 | 53.54 | 3.57 | 0.83 | 195.05 | 31.82 | 31.45 | 3.89 | 0.48 | 26.69 | 0.40 |
| thrb | 59.91 | 16 | 47.23 | 46.99 | 1.66 | 0.79 | 1174.91 | 25.36 | 25.4 | 1.65 | 0.42 | 5.9 | 0.10 |
| try1 | 58.56 | 21 | 58.11 | 57.95 | 0.6 | 0.99 | 1123.07 | 44.76 | 44.62 | 1.45 | 0.76 | 5.57 | 0.10 |
| tryb1 | 52.41 | 16 | 44.35 | 43.59 | 2.29 | 0.85 | 256.36 | 25.53 | 24.95 | 2.87 | 0.49 | 5.38 | 0.10 |
| tysy | 62.61 | 36 | 35.39 | 33.72 | 3.66 | 0.57 | 272.54 | 19.96 | 19.87 | 3.54 | 0.32 | 12.7 | 0.20 |
| urok | 61.56 | 28 | 60.94 | 59.83 | 1.77 | 0.99 | 419.33 | 57.86 | 57.2 | 1.92 | 0.94 | 25.24 | 0.41 |
| vgfr2 | 57.28 | 17 | 28.51 | 28.16 | 1.8 | 0.50 | 831.42 | 15.62 | 15.67 | 1.78 | 0.27 | 8.43 | 0.15 |
| wee1 | 61.16 | 38 | 61.16 | 61.15 | 0.09 | 1.00 | 222.52 | 61.16 | 61.12 | 0.24 | 1.00 | 40.77 | 0.67 |
| xiap | 52.3 | 44 | 52.3 | 52 | 0.82 | 1.00 | 194.12 | 49.34 | 48.61 | 1.98 | 0.94 | 27.63 | 0.53 |
| Mean | 62.24 | 26.46 | 40.40 | 39.59 | 2.72 | 0.65 | 545.77 | 25.09 | 24.92 | 2.74 | 0.41 | 12.02 | 0.20 |

[(a)] Target identification on the DUD-E database (http://dude.docking.org/)

[(b)] EF at 1% of screened data for a perfect ranking of the active compounds on the ranked list

[(c)] Number of aggregated scoring functions

[(d)] EF of the best performing VS strategy

[(e)] Mean EF on 1000 bootstrap simulations of the best performing VS strategy

[(f)] Standard deviation of the bootstrap cross-validation procedure

[(g)] Fraction of the maximum possible EF that the VS method achieves

[(h)] Run time for each consensus scoring approach

Table TS6. VS performance on external validation experiments. Enrichment is computed as BEDROC with α is set to 160.9. Results are presented as the average BEDROC over the 100 training/external partitions of each target

| Target[(a)] | BEDROC | | | |
|---|---|---|---|---|
| | Train[(b)] | Std (Train)[(c)] | Ext.[(d)] | Std (Ext.)[(e)] |
| aa2ar | 0.73 | 0.01 | 0.71 | 0.05 |
| abl1 | 0.60 | 0.02 | 0.57 | 0.07 |
| ace | 0.52 | 0.02 | 0.48 | 0.06 |
| aces | 0.65 | 0.01 | 0.62 | 0.06 |
| ada | 0.76 | 0.02 | 0.64 | 0.10 |
| ada17 | 0.78 | 0.01 | 0.77 | 0.04 |
| adrb1 | 0.48 | 0.02 | 0.41 | 0.07 |
| adrb2 | 0.44 | 0.02 | 0.35 | 0.07 |
| akt1 | 0.69 | 0.01 | 0.63 | 0.06 |
| akt2 | 0.64 | 0.02 | 0.59 | 0.11 |
| aldr | 0.60 | 0.03 | 0.53 | 0.10 |
| ampc | 0.67 | 0.04 | 0.52 | 0.19 |
| andr | 0.54 | 0.02 | 0.50 | 0.07 |
| aofb | 0.41 | 0.03 | 0.26 | 0.10 |
| bace1 | 0.33 | 0.02 | 0.24 | 0.07 |
| braf | 0.61 | 0.02 | 0.55 | 0.09 |
| cah2 | 0.52 | 0.01 | 0.49 | 0.05 |
| casp3 | 0.38 | 0.03 | 0.30 | 0.09 |
| cdk2 | 0.48 | 0.01 | 0.43 | 0.06 |
| comt | 0.99 | 0.01 | 0.96 | 0.04 |
| cp2c9 | 0.27 | 0.02 | 0.09 | 0.07 |
| cp3a4 | 0.26 | 0.02 | 0.14 | 0.07 |
| csf1r | 0.51 | 0.03 | 0.44 | 0.09 |
| cxcr4 | 0.73 | 0.03 | 0.51 | 0.16 |
| def | 0.91 | 0.01 | 0.88 | 0.05 |

| | | | | |
|---|---|---|---|---|
| dhi1 | 0.35 | 0.02 | 0.30 | 0.08 |
| dpp4 | 0.64 | 0.01 | 0.63 | 0.05 |
| drd3 | 0.43 | 0.01 | 0.41 | 0.05 |
| dyr | 0.75 | 0.01 | 0.72 | 0.06 |
| egfr | 0.75 | 0.01 | 0.73 | 0.04 |
| esr1 | 0.76 | 0.01 | 0.74 | 0.05 |
| esr2 | 0.74 | 0.01 | 0.72 | 0.05 |
| fa10 | 0.72 | 0.01 | 0.71 | 0.04 |
| fa7 | 0.95 | 0.01 | 0.93 | 0.03 |
| fabp4 | 0.77 | 0.04 | 0.60 | 0.16 |
| fak1 | 0.70 | 0.03 | 0.58 | 0.12 |
| fgfr1 | 0.56 | 0.03 | 0.47 | 0.11 |
| fkb1a | 0.71 | 0.03 | 0.64 | 0.11 |
| fnta | 0.22 | 0.01 | 0.19 | 0.04 |
| fpps | 0.91 | 0.01 | 0.89 | 0.05 |
| gcr | 0.48 | 0.02 | 0.41 | 0.08 |
| glcm | 0.85 | 0.02 | 0.72 | 0.13 |
| gria2 | 0.70 | 0.02 | 0.65 | 0.07 |
| grik1 | 0.84 | 0.02 | 0.77 | 0.08 |
| hdac2 | 0.74 | 0.02 | 0.70 | 0.07 |
| hdac8 | 0.90 | 0.01 | 0.88 | 0.04 |
| hivint | 0.33 | 0.02 | 0.15 | 0.07 |
| hivpr | 0.30 | 0.01 | 0.27 | 0.04 |
| hivrt | 0.44 | 0.02 | 0.37 | 0.06 |
| hmdh | 0.83 | 0.01 | 0.80 | 0.07 |
| hs90a | 0.77 | 0.02 | 0.69 | 0.11 |
| hxk4 | 0.75 | 0.02 | 0.67 | 0.10 |
| igf1r | 0.55 | 0.02 | 0.43 | 0.10 |
| inha | 0.73 | 0.04 | 0.43 | 0.18 |
| ital | 0.45 | 0.03 | 0.39 | 0.12 |
| jak2 | 0.80 | 0.02 | 0.72 | 0.09 |

| | | | | |
|---|---|---|---|---|
| kif11 | 0.83 | 0.02 | 0.78 | 0.07 |
| kit | 0.38 | 0.02 | 0.30 | 0.09 |
| kith | 0.96 | 0.01 | 0.92 | 0.05 |
| kpcb | 0.74 | 0.02 | 0.69 | 0.08 |
| lck | 0.61 | 0.01 | 0.60 | 0.05 |
| lkha4 | 0.61 | 0.02 | 0.48 | 0.10 |
| mapk2 | 0.92 | 0.01 | 0.87 | 0.06 |
| mcr | 0.53 | 0.04 | 0.43 | 0.14 |
| met | 0.67 | 0.02 | 0.64 | 0.08 |
| mk01 | 0.63 | 0.03 | 0.43 | 0.15 |
| mk10 | 0.57 | 0.02 | 0.41 | 0.11 |
| mk14 | 0.41 | 0.01 | 0.39 | 0.05 |
| mmp13 | 0.68 | 0.01 | 0.66 | 0.04 |
| mp2k1 | 0.40 | 0.02 | 0.31 | 0.10 |
| nos1 | 0.68 | 0.02 | 0.61 | 0.10 |
| nram | 0.95 | 0.01 | 0.93 | 0.04 |
| pa2ga | 0.74 | 0.02 | 0.66 | 0.12 |
| parp1 | 0.90 | 0.01 | 0.90 | 0.02 |
| pde5a | 0.57 | 0.01 | 0.54 | 0.05 |
| pgh1 | 0.41 | 0.02 | 0.33 | 0.09 |
| pgh2 | 0.63 | 0.01 | 0.60 | 0.05 |
| plk1 | 0.71 | 0.02 | 0.61 | 0.10 |
| pnph | 0.91 | 0.01 | 0.87 | 0.05 |
| ppara | 0.41 | 0.04 | 0.36 | 0.08 |
| ppard | 0.46 | 0.02 | 0.40 | 0.09 |
| pparg | 0.46 | 0.02 | 0.44 | 0.06 |
| prgr | 0.76 | 0.01 | 0.72 | 0.05 |
| ptn1 | 0.71 | 0.02 | 0.62 | 0.10 |
| pur2 | 0.97 | 0.01 | 0.92 | 0.05 |
| pygm | 0.43 | 0.06 | 0.23 | 0.13 |
| pyrd | 0.82 | 0.02 | 0.76 | 0.07 |

| | | | | |
|---|---|---|---|---|
| reni | 0.67 | 0.02 | 0.61 | 0.09 |
| rock1 | 0.69 | 0.02 | 0.57 | 0.11 |
| rxra | 0.95 | 0.01 | 0.94 | 0.03 |
| sahh | 0.98 | 0.01 | 0.93 | 0.06 |
| src | 0.53 | 0.01 | 0.52 | 0.05 |
| tgfr1 | 0.85 | 0.01 | 0.83 | 0.06 |
| thb | 0.81 | 0.02 | 0.74 | 0.08 |
| thrb | 0.76 | 0.01 | 0.75 | 0.04 |
| try1 | 0.92 | 0.01 | 0.91 | 0.02 |
| tryb1 | 0.79 | 0.01 | 0.76 | 0.07 |
| tysy | 0.63 | 0.02 | 0.53 | 0.11 |
| urok | 0.93 | 0.01 | 0.91 | 0.03 |
| vgfr2 | 0.48 | 0.02 | 0.45 | 0.06 |
| wee1 | 1.00 | 0.00 | 0.98 | 0.01 |
| xiap | 0.96 | 0.01 | 0.94 | 0.03 |

[a] Target identification on the DUD-E database (http://dude.docking.org/)

[b] Mean BEDROC on the training set over 100 data splits

[c] Standard deviation of BEDROC on the training set over 100 data splits

[d] Mean BEDROC on the external set over 100 data splits

[e] Standard deviation of BEDROC on the external set over 100 data splits
